# The sensory representation of causally controlled objects

**DOI:** 10.1101/786467

**Authors:** Kelly B. Clancy, Thomas D. Mrsic-Flogel

## Abstract

Intentional control over external objects is informed by our sensory experience of them. To study how causal relationships are learned and effected, we devised a brain machine interface (BMI) task utilising wide-field calcium signals. Mice learned to entrain activity patterns in arbitrary pairs of cortical regions to guide a visual cursor to a target location for reward. Brain areas that were normally correlated could be rapidly reconfigured to exert control over the cursor in a sensory feedback-dependent manner. Higher visual cortex was more engaged when expert but not naïve animals controlled the cursor. Individual neurons in higher visual cortex responded more strongly to the cursor when mice controlled it than when they passively viewed it, with the greatest response boosting as the cursor approached target location. Thus, representations of causally-controlled objects are sensitive to intention and proximity to the subject’s goal, potentially strengthening sensory feedback to allow more fluent control.

## Introduction

How does the brain infer a causal relationship between its activity and the sensed world, and how does this affect the sensory encoding of controlled external objects? Actions and perceptions reciprocally affect one another in a mutual dialog (Dewey, 1896). The sense of causal control can be operationalised to inferring a causal relationship between a subject’s internally generated actions or activity, and their outcome in the external world (Haggard, 2017). In motor learning, for example, the relationship between an action and its outcome can be learned and re-learned throughout adulthood as animals acquire new motor skills. Brain machine interfaces (BMI) are a method for investigating how subjects learn arbitrary action-outcome relationships (Fetz, 1969; Bakay and Kennedy, 1998; Nicolelis, 2001; Donoghue, 2002; Carmena et al., 2003; Sitaram et al., 2017). When learning to control a BMI, rodents have been found to employ the same mechanisms as implicated in motor learning (Koralek et al., 2012; Neely et al., 2018). But unlike motor learning, wherein animals learn a task and researchers must search for correlates of the behaviour in patterns of neural activity, BMIs allow the experimenter to precisely control sensory feedback, as well as prescribe the requisite activity patterns necessary for successful task execution, which can then be changed day to day. Thus, animals learning neuroprosthetic control of external objects must engage in continuous self-monitoring to assess the contingency between their neural activity and its outcome, preventing them from executing a habitual or fixed motor pattern, and encouraging animals to learn arbitrary new sensorimotor mappings on the fly.

A key aspect of this self-monitoring is the sensory feedback from the object being controlled by the agent. Yet little is known about how causally controlled objects are represented in the brain. Studies have implicated parietal cortex in intention, as well as in the subjective assessment of agency over outcome. In human subjects, disrupting activity in parietal cortex with TMS temporarily ablates self-reported agency (Chambon et al., 2015). Parietal activity has been found to be involved in representing task rules, the value of competing actions, and visually-guided real-time motor plan updating: both in humans (Pisella et al., 2000; Kahnt et al., 2014; Wisniewski et al., 2015; Zapparoli et al., 2020) and non-human primates (Andersen et al., 1997; Sugrue et al., 2004). Motor plans can be decoded from parietal activity, and its responses are task-, expectation-, and goal-dependent, in humans, (Rushworth et al., 2001; Desmurget et al., 2009; Aflalo et al., 2015), non-human primates (Mountcastle et al., 1975; Gnadt and Andersen, 1988; Lacquaniti et al., 1995; Johnson et al., 1996; Churchland et al., 2008), and rodents (Licata et al., 2017; Pho et al., 2018; Mohan et al., 2018, 2019). All of this evidence suggests that, across multiple species, parietal cortex plays a role in intentional, goal directed behaviours (Rizzolatti et al., 1997; Andersen and Buneo, 2002; Andersen and Cui, 2009). However, previous studies of the role of parietal cortex in intention have not examined how causal control affects sensory representations across different sensorimotor contingencies.

To address this, we devised a mouse model of adaptive causal control. Animals learned to guide a visual feedback cursor to a target location in order to obtain a reward using activity in experimenter-defined cortical areas, recorded with widefield imaging. This had the added benefit of acting as an unbiased screen to identify dorsal cortical areas involved in learning the task. We found that higher visual areas, including anteromedial cortex, AM, were more engaged when expert animals controlled the BMI. These higher areas are considered by some to be a putative homologue of parietal cortex in mice (Harvey et al., 2012; Licata et al., 2017; Mohan et al., 2018; Pho et al., 2018; Lyamzin and Benucci, 2019). In order to gain insight into what this task-related activity looks like on an individual neuron level, we targeted single-cell recordings to the functionally-identified task-relevant region AM, and found that the visual cursor elicited larger responses when an animal was controlling it in a closed-loop configuration than when passively viewing it in an open-loop configuration (Bagur et al., 2018). Responses were highest when the cursor was closest to the target zone, and were sensitive to the cursor’s instantaneous trajectory: they were greater when the cursor was moving towards the target than away from it. Thus, the sensory representation of the visual object was sensitive to the subject’s intention, as well as its perception of the object’s instantaneous trajectory with respect to its goal. Given that animals had to relearn a changing sensorimotor contingency on the fly, we surmised that the heightened sensory representation of the cursor at the target position might serve to strengthen the signal to adjudicating areas for informing fluent control over external objects. Indeed, the neural activity in AM during the task condition carried significantly more information about the cursor identity than during the passive playback condition.

## Results

### Goal-directed control of a visual cursor using areal signals

In order to investigate how causal control over external objects is effected and encoded in mammalian cortex, we trained mice to control a visual feedback cursor using real-time calcium signals recorded with wide-field imaging (largely reflecting the summed spiking activity of local cells (Makino et al., 2017; Clancy et al., 2019). We imaged the dorsal cortex in transgenic mice expressing the calcium indicator GCamp6s in CaMKII+ pyramidal neurons (Wekselblatt et al., 2016), assigning two small frontal regions to control a visual feedback cursor (Figure 1A-B, Supplemental Video 1-3), similar to a task described previously (Clancy et al., 2014; Koralek et al., 2012). Animals were head-fixed under the widefield microscope, and free to run on a styrofoam wheel. The animal’s goal was to bring a visual cursor (a copy of which was presented to both eyes on two separate monitors flanking each side of the mouse) to a target position in the centre of its visual field (Figure 1C). Animals could achieve this by increasing activity in control region 1 (R1) relative to control region 2 (R2). If activity in R2 was greater than R1, the cursor moved towards the back of the animal’s visual field. Thus, in this design, animals could not hit the target simply by generally increasing or decreasing activity across cortex, but had to differentially modulate the activity of these experimenter-specified control regions (Figure 1D-F; Supp. Figure 1A-B). These control regions were usually placed over ipsilateral motor areas and were changed from day to day (Figure 1B, Supplemental Table 1). Control regions were deliberately small (~0.1 mm^2^) as we reasoned that it would be easier for animals to control a smaller area, but this was not systematically tested. We avoided using anterior lateral motor cortex (ALM) in the control regions, to minimise the effect of licking.

**Figure 1.**
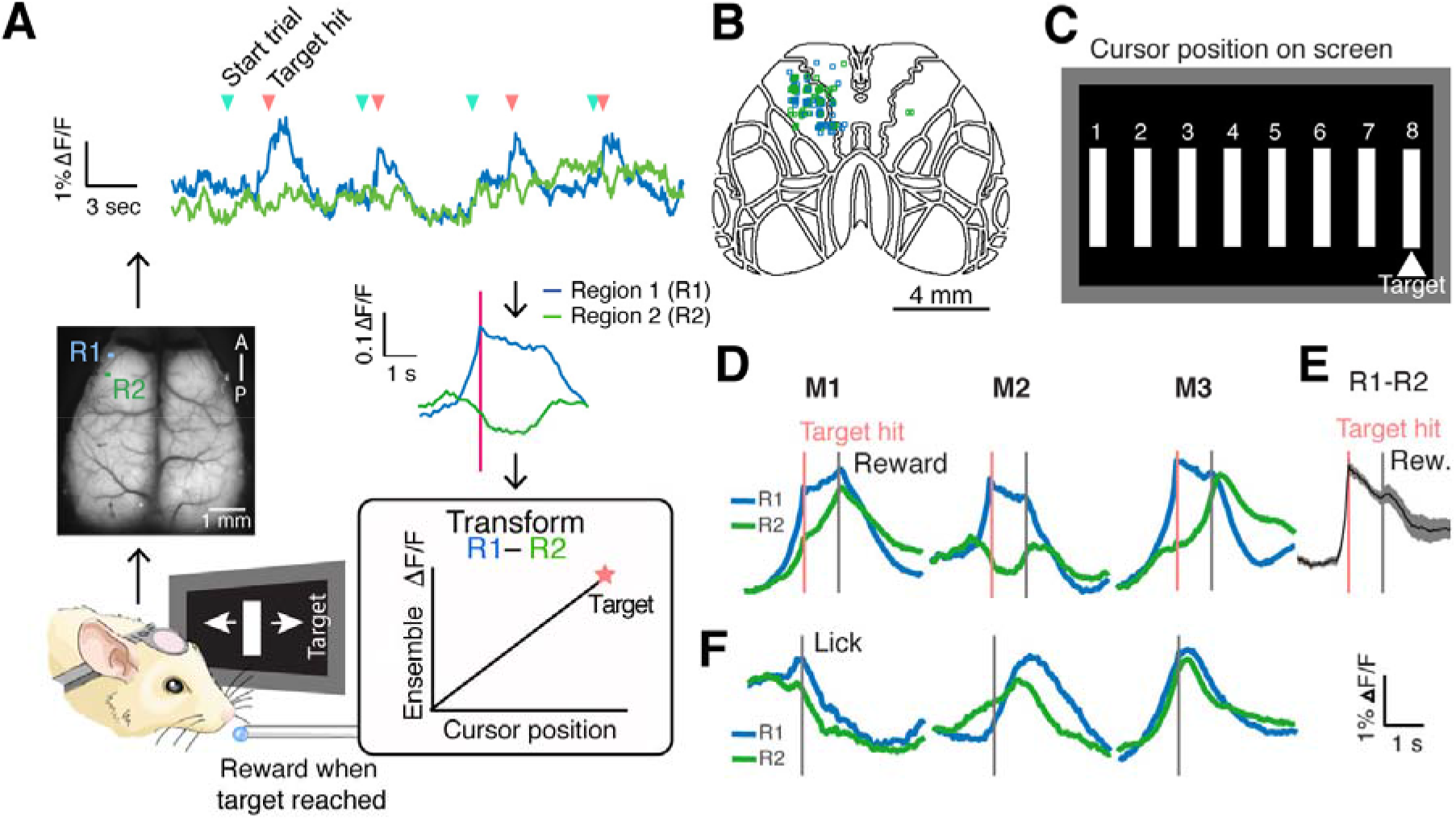
A widefield-imaging based brain machine interface. **a.** Task schematic. Clockwise starting from cartoon mouse: widefield signals were imaged from head-fixed animals in real time and transmuted into the position of a visual cursor. Two small regions (R1 and R2) were used for controlling the cursor, and activity recorded from these areas was fed into a decoder such that their activity opposed one another. Example dF/F for the two regions is shown at top, with blue arrows denoting trial starts, and pink arrows denoting target hits. Activity averaged around hits for one example animal, one day, shown for R1 and R2. Animals had to increase activity in R1 relative to R2 to bring the cursor to a rewarded position at the centre of their visual field, at which point they could collect a reward after a 1 second delay. **B.** Positions of control ROIs (R1 in blue, R2 in green) for all 7 animals over the course of training (averaging 15 days each), superimposed on the Allen Brain Atlas (totalling 104 pairs). **C.** Feedback schematic: the cursor could take one of 8 potential positions on screen, with position 8, the target, rewarded. **D.** ΔF/F in control regions triggered around hits for three example animals on one day of training, indicating different strategies animals use to achieve reward. Pink line indicates the time of target hit, grey line indicates reward delivery. **E.** Activity in R1 subtracted by activity in R2, averaged around target hits for all mice on a day of training (n=7 mice, shading represents s.e.m.). **F.** Animals could not control cursor using lick alone. ΔF/F triggered around lick bouts in spontaneous activity for the same three example animals on one day of training, indicating animals could not achieve the differential activation of R1 and R2 using lick alone. Grey line indicates time of lick.

The visual feedback cursor could take one of eight positions on the monitors (Figure 1C), and the cursor had to be held at the target for 300 milliseconds to count as a hit. When animals succeeded in holding the cursor at the target position, a soya milk reward was delivered after a one second delay. If animals failed to bring the cursor to the target position within 30 seconds, the trial was considered a miss, and a white noise miss-cue was played, followed by a brief time out. Chance performance was assessed using spontaneous activity recorded before the task began, and represents the estimated hit rate that the animal would achieve using spontaneous fluctuations of neural signals alone. Animals improved their performance over training (Figure 2A, n = 7 mice), and took less time to reach a criterion performance of 50% hit trials over days (Figure 2B).

**Figure 2.**
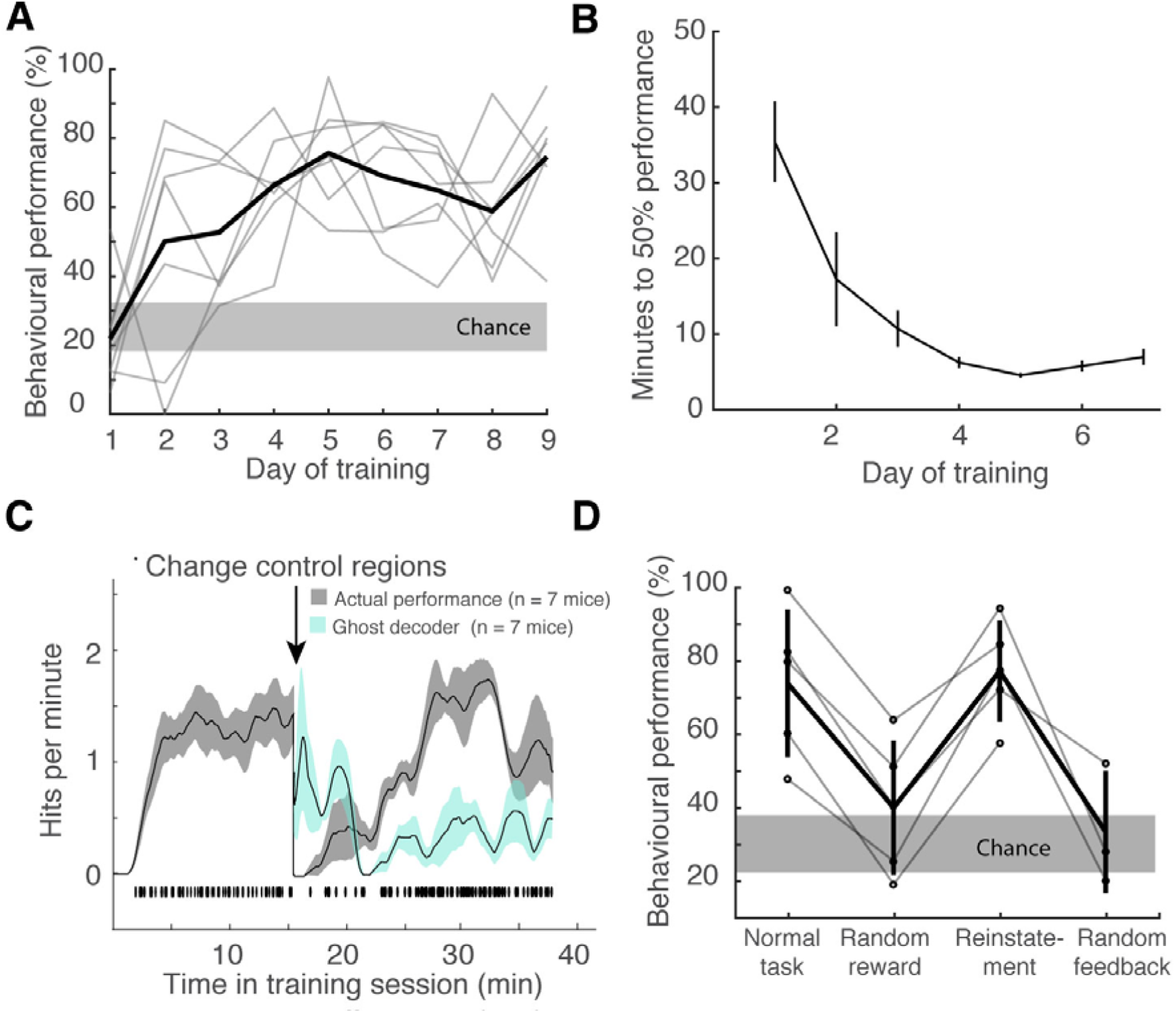
Animals learn to control a visual cursor using areal neural activity. **A.** Behavioural performance (percentage of trials the animal successfully reached the target within the 30 second trial window) increased above chance over the course of days. Shaded region denotes chance s.e.m., assessed as how often spontaneous activity would have achieve hits, averaged across 7 mice. **B.** Animals achieve 50% performance faster over the course of days of training, calculated by taking a moving average of the number of times the animal successfully reached the target per trial. **C.** Average hits per minute (grey) increased over the course of a training session, and recovered within minutes when control regions were changed (n = 7 mice on day six of training, shading denotes 95% confidence interval, see Supplemental Table 1). Individual hit times, pooled for all animals, shown below rate curves. At chance baseline, animals should perform around 0.3 hits/minute (here, animals started at slightly worse than chance, likely due to the fact that some of the control regions positions had been changed from the previous day). After R1 and R2 were swapped mid-session, the estimate of how well the animals would have done using the original control regions is shown in teal: the previous (“ghost”) decoder’s performance starts near the animal’s previous hit rate, suggesting they persist with their original strategy, but this drops off as animals discover the new rewarded contingency and recover their performance. **D.** Performance dropped to near chance when rewards were given randomly, but recovered when target and reward was again coupled on a subsequent day (n = 5 mice). Performance also dropped to chance when the visual feedback cursor was uncoupled from neural activity and presented at random (n = 3 mice), even though animals could still achieve reward using the previously learned activity pattern.

Animals could perform the task without overt movements, licking, or eye saccades (Supp. Figure 1C-E). Several recent papers have shown that many motor activities can influence cortical activity (Stringer et al., 2019; Musall et al., 2019; Orsolic et al., 2019; Salkoff et al., 2020), raising the possibility that mice may adopt a motor strategy to control the activation of cortical regions used for BMI. Our analyses show that animals could perform the task without gross overt movements that were detected by monitoring licking, eye saccades or running speed (Supp. Figure 1C-E), though it is possible that subtler movement were not detected. This is a question, common to all BMI studies, that is ultimately unanswerable without full recordings from every muscle in the body. Here we did not take full body videos of the animals performing the task, but in future work including video monitoring can offer insights on subtler motor outputs. Nevertheless, animals in this study could discover arbitrary activation patterns of different brain areas within and between training sessions, and use these activations to control a visual cursor in a manner dependent on sensory feedback: that is, they could flexibly work out the arbitrary coupling of action patterns (whether purely confined to the brain, or involving peripheral muscle contractions) with sensory feedback in order to achieve a goal. The flexibility with which the animals could rapidly readjust their control of arbitrary brain areas suggests some degree of adaptability in decorrelating otherwise highly correlated brain areas (see below).

Control regions could be changed from day to day or within a session: hit rates improved over time within a session, and recovered after control regions were changed, indicative of learning (Figure 2C, Supp. Figure 2A-B). After recovery, the animals’ hit rate before and after this control region switch were not statistically different (Supp. Figure 2C). The decoder transformation for both the pre- and post-switch conditions was calculated using the same spontaneous baseline taken before the training session. To test if the behaviour was goal-directed, we dissociated the reward from the target position. On day 8 of training, animals were allowed to perform the task as usual, but the training session was constrained to 30 minutes. Thereafter, the visual feedback was coupled to the animal’s neural activity as before, but rewards were given randomly, at the same rate as an expert animal engaged in the task (~1.3 hits/minute). During this random reward condition, the animals’ target hit rate dropped to chance, indicating that the behaviour was goal-oriented (Figure 2D). The fluorescence signals representing the difference between R1 and R2 increased over the normal training session, indicative of increased efficacy of control, and decreased when reward was randomised, again suggesting that animals were effecting the requisite neural patterns in a goal-directed manner (Supp. Figure 2D-E). Animals were able to recover their performance following reinstatement of the normal task on the next day of training (Figure 2D). For a subset of animals, the visual feedback was then randomised (referred to as the random feedback condition): the visual cursor was presented at random positions, though animals could still achieve the target with the appropriate neural activity patterns independent of cursor position. The animals’ ability to bring the cursor to the target dropped to chance levels without meaningful visual feedback (Figure 2D). Activity leading to hits did not resemble activity induced by licking around reward collection, as evidenced in both the normal task and random-reward conditions (Supp. Figure 2F). Altogether, this suggests that animals performed the task in a flexible, goal-directed, visual feedback-dependent manner.

### Exploration and exploitation in neural activity space

As control regions were changed day-to-day, the activity patterns necessary for successful BMI control had to be re-learned each session. Example fluorescence traces from control regions indicate the areas were initially highly spontaneously correlated (Figure 3A, top trace; Supplemental Figure 1A-B; Supplemental Video 3). Early in the training session, hits were preceded by diverse activity patterns (Figure 3A, middle trace). By late in the training session, the activity patterns leading to hits were more consistent (Figure 3A, bottom trace). Animals found different ways to achieve this consistency, sometimes by sweeping activity through R1 towards R2, or by depressing R2 while activating R1 (Figure 1D, Supp. Videos 1-3).

**Figure 3.**
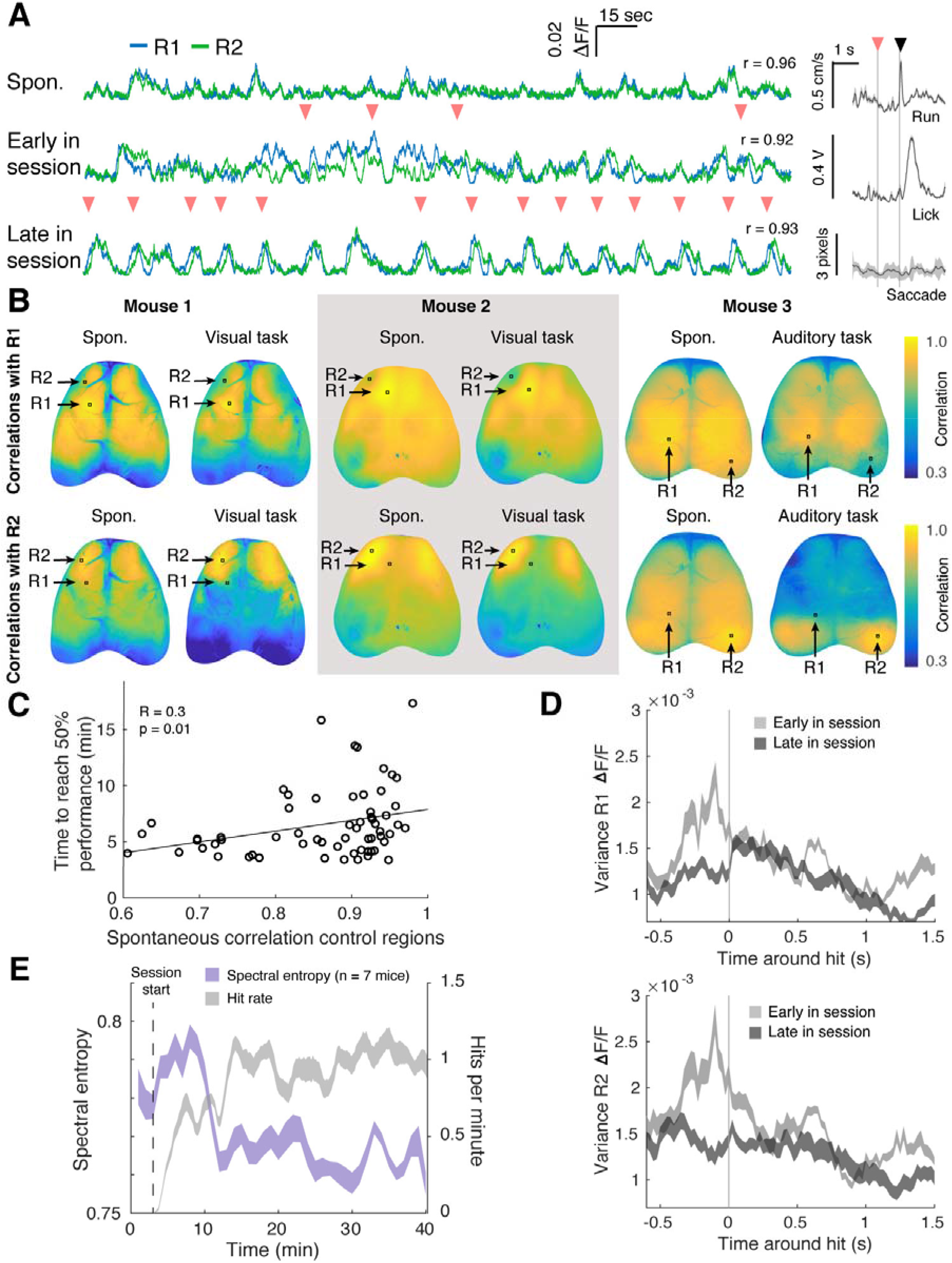
Exploration and exploitation of neural activity patterns. **A.** Areal signals were highly spontaneously correlated before the training session (top trace). Animals explored different activation patterns early in the training session (middle trace) in order to discover and exploit successful patterns by the end of the session (bottom trace). Pink arrows denote target hits. Pearson’s correlation between R1 and R2 indicated on the right of each trace. Right: run, lick and saccade averaged around hits for this training session, grey shading denotes 95% confidence interval around mean. Pink arrow denotes target hits, black arrow denotes reward delivery. **B.** Correlation map across cortex for 3 animals using activity in R1 (top row) and R2 (bottom row) as seeds, during spontaneous activity and during the BMI task. Mouse 3 was part of a separate cohort trained on an auditory task (see Methods). Animals could decorrelate normally correlated areas for task performance. **C.** Animals took longer to reach criterion performance (50% hits/attempt) if control regions were highly spontaneously correlated (data from 7 mice, 9 days of training starting on day 4). **D.** The average variance of activity for R1 (top panel) and R2 (bottom panel) was greater around hits early in a training session than late in the session (n = 7 mice, day 8 of training, shading indicates 95% confidence interval around mean) indicating that mice honed in on more reliable and reproducible control strategies over the course of a single training session. **E.** Early in the session, neural activity around the control regions had high spectral entropy (a proxy of signal randomness) as animals used stochastic bursts of activity to explore the neural patterns that would yield reward. By late in the session, animals had discovered a successful activity pattern to exploit, and spectral entropy in control area activity decreased. Shaded area indicates 95% confidence interval around mean.

In order to achieve the prescribed activity pattern, animals had to functionally decorrelate the two control regions, which were usually highly spontaneously correlated. Indeed, activity in dorsal cortex was globally correlated in animals pre-task, as indicated by correlation maps using R1 and R2 as seed pixels (Figure 3B; Supp. Figure 1B). Although activity in dorsal cortex was globally correlated in animals pre-task, correlations between these areas decreased during task performance (Figure 3B, Supp. Figure 3B), as did the correlations between the control regions and primary visual cortex, V1, (Supp. Figure 3C). Correlations between the control regions and primary somatosensory cortex, S1, as well as between S1 and V1 were unchanged between spontaneous activity ans the task (Supp. Figure 3C-D). Only periods of task performance, and not reward collection or inter-trial waiting periods, were included in these analyses. Animals could arbitrarily decorrelate different regions: in a separate cohort of animals trained using auditory feedback instead of the visual feedback cursor, animals could also decorrelate visual control areas (Figure 3B, rightmost panel; Supp. Figure 4). Interestingly, the task-induced correlation patterns were invariably bilateral, even when control regions were ipsilateral to each other. Not every training session resulted in the kind of decorrelated maps evident in Figure 3B; reflecting that some areas are harder for the animals to decorrelate than others: animals took longer to reach criterion 50% performance when control regions were spontaneously highly correlated (Figure 3C). The distance between control regions did not affect the time animals required to reach 50% performance (Supp. Figure 3A), suggesting that areas that were the most spontaneously correlated, irrespective of their distance from one another, were the hardest to use to perform the task.

The variance in R1 and R2 activity peaked around hits early in a training session as animals explored strategies that would yield reward, as has been shown previously for the activity of individual cells learning to control a BMI device using a fixed decoder (Zacksenhouse et al., 2007; Athalye et al., 2017). This variance decreased later in the session as animals discovered and exploited reliable, reproducible strategies (Figure 3D). Early in the session, the spectral entropy of activity (a measure of the spectral power distribution of a signal, and a proxy for its complexity, see Methods) around the control regions increased as animals explored activity patterns that would yield reward, then dramatically dropped as they discovered successful patterns and reliably exploited them (Figure 3E). At the start of a session, or when control regions were changed, animals faced with uncertain task rules ‘explored’ activity space through stochastic bursting, and gradually switched to effecting stereotyped patterns they could reliably exploit after having probed the rules of their environment (Tervo et al., 2014).

### Expert performance correlated with increased activity in higher visual areas

We sought to determine what cortical areas were most active during task performance, and how this changed over learning. On the first day of training, primary visual cortex was most active during the task, but as animals became expert in the task, higher visual areas were recruited (Figure 4A-C): in particular, anteromedial cortex (AM), posteriomedial cortex (PM), and rostrolateral visual cortex (RL), similar to previous studies of learning visually-guided tasks in mice (Wekselblatt et al., 2016; Orsolic et al., 2019). Areas AM and RL are considered by some to be parietal cortex homologues in mouse cortex, though there is no widespread agreement on this point (Harvey et al., 2012; Licata et al., 2017; Mohan et al., 2018; Pho et al., 2018; Lyamzin and Benucci, 2019). After the final day of training, animals were shown playback of the cursor positions using their previous task performance (hereafter referred to as the passive playback condition). Activity in these higher areas was not evident in animals passively watching the cursor, suggesting their recruitment was specific to goal-oriented task engagement (Figure 4). A separate cohort of animals trained using an auditory feedback cursor had variable task-active areas (Supp. Figure 4A, n = 4 mice), but, as with the visual task, higher activity was also evident in RL, which is a multimodal sensory associative area with neurons responsive to visual, touch and auditory stimuli (Mohan et al., 2018).

**Figure 4:**
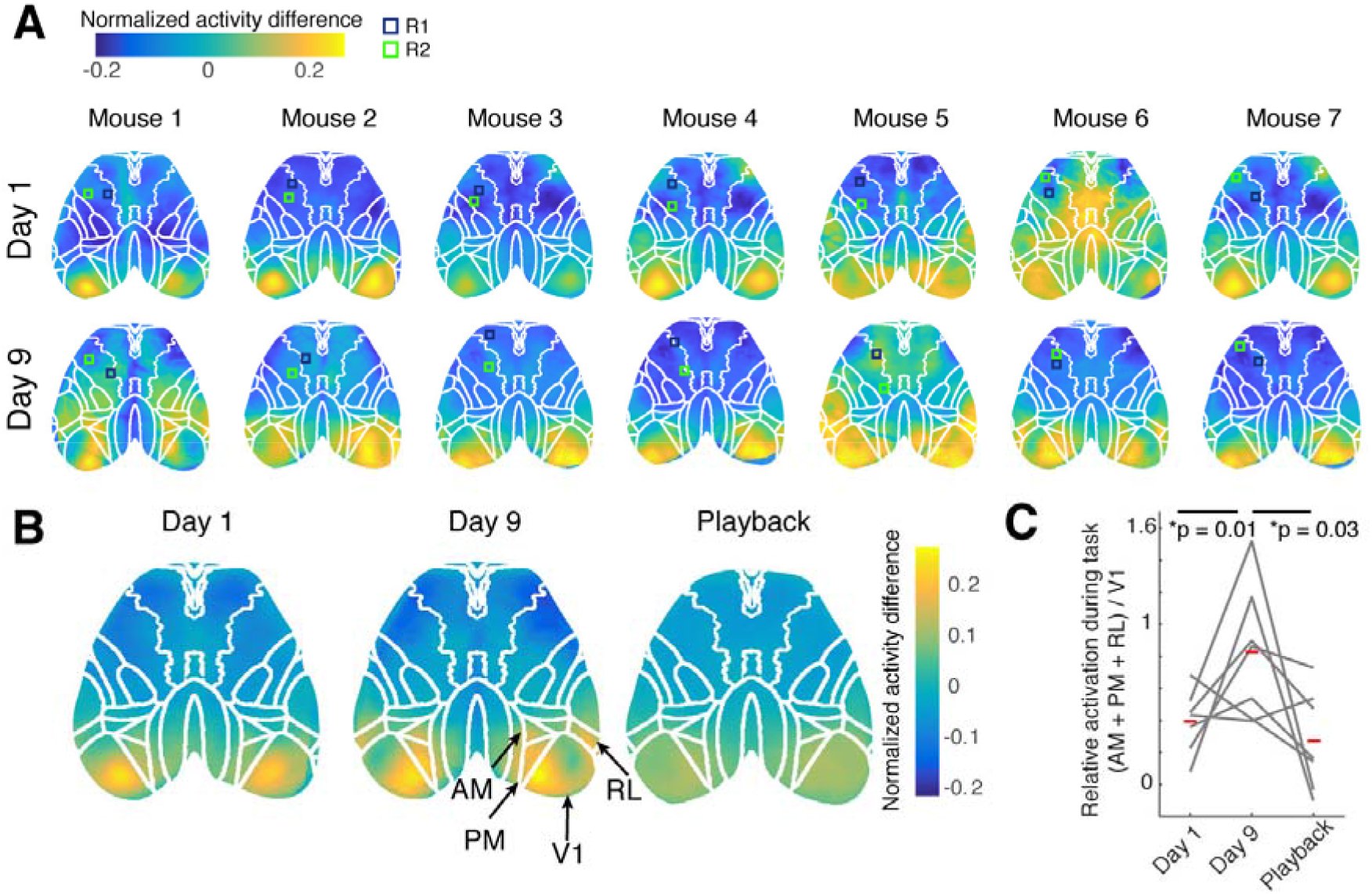
Higher visual areas were more active during expert task performance. **A.** Activation maps for individual animals on day 1 of training (top row), when animals were naïve, versus day 9 of training (bottom row), when animals performed the task expertly, calculated using the normalised activity difference for task-on minus task-off periods. Each map has been registered to the Allen Brain Atlas (overlaid) using stereotaxic marks. Control regions are shown as slightly larger than they actually were for better visibility. **B.** Activation maps during task performance on day 1, day 9, and during passive playback of a previous session (representing the normalised activity difference for task-on minus task-off periods). Each map has been registered to the Allen Brain Atlas by stereotaxic marks and then averaged across 7 mice. **C.** The relative ratio of task-activation in higher visual areas versus V1 increased over training. When animals passively viewed playback of the same session, higher visual areas were less active. Red bars indicate mean ratios (n=7 mice, paired t-test, Bonferroni corrected).

### Population tuning of neurons shifted towards target position

Having identified brain areas implicated in BMI control, we recorded spiking from individual cells while animals performed the task, in order to investigate the task-dependent increase in calcium signals with cellular resolution. We chose to record from functionally identified area AM, due to its recruitment over learning, and used multi-channel silicon probes to record spikes from individual neurons while simultaneously imaging the rest of dorsal cortex (Figure 5A, see Methods, (Xiao et al., 2017; Clancy et al., 2019; Barson et al., 2018; Peters et al., 2019)). We obtained 16-49 single units per recording, spanning all cortical layers. Units could be classified as regular-spiking (RS) or fast-spiking, putative interneurons (FS) depending on spike width (see Methods, Figure 5A). We recorded from 131 units in 7 mice performing the task (example recording shown in Figure 5B), and 128 units from the same animals passively viewing a playback of the previous training session’s cursor positions. Due to electrode shift over the recording sessions, we do not have recordings of the same units across both conditions, thus our analyses are limited to population response differences. During the task, population firing was significantly increased for cursor positions closer to the target, and this was true both of FS and RS units (Figure 5C-E, Supp. Figure 5A-C). The target-dependent increased spiking could not be explained by reward expectation, as a subset of animals were given reward at the target position during passive playback as well (Supp. Figure 5C).

**Figure 5.**
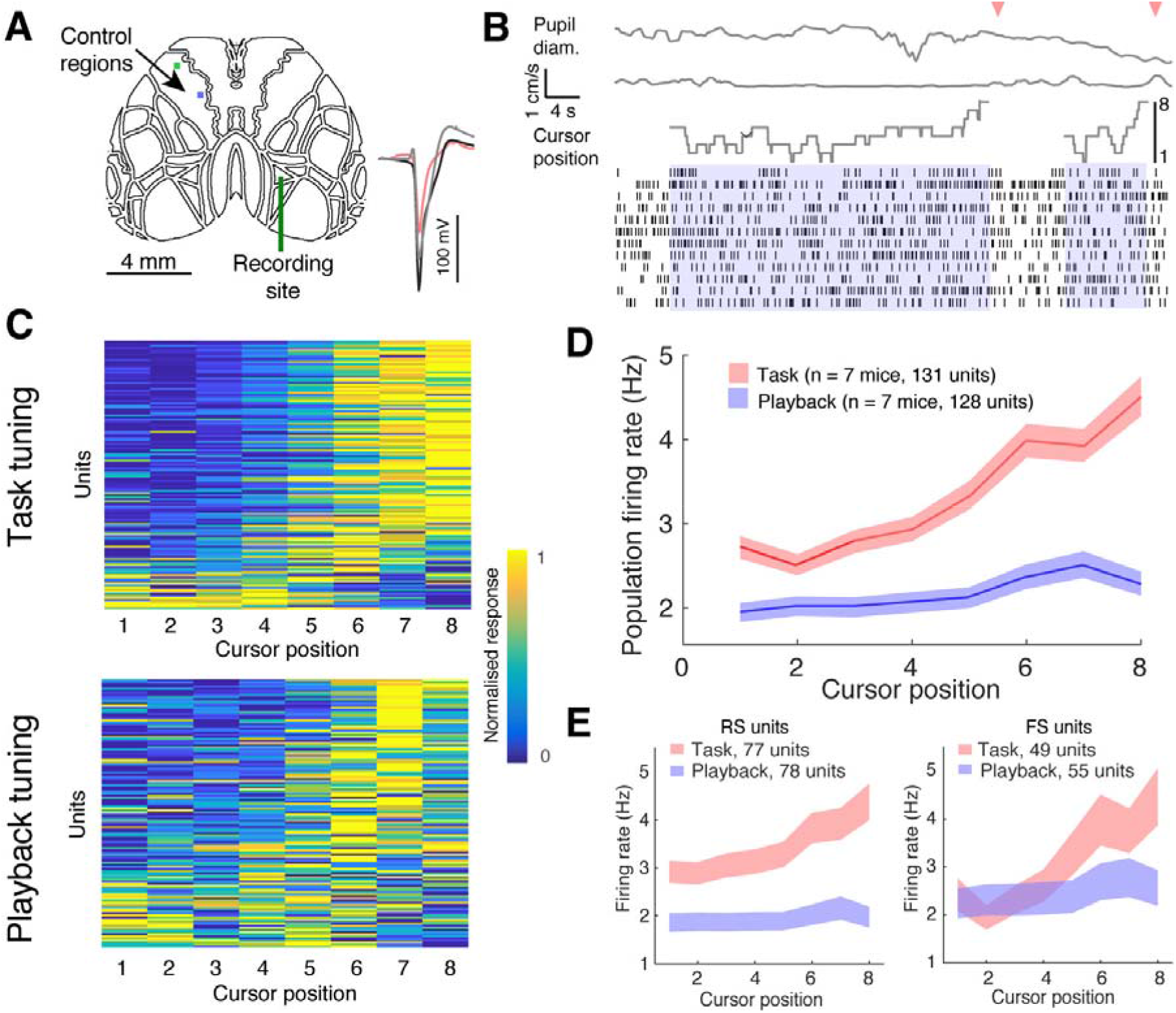
Cursor tuning shifts to target location. **A.** Electrophysiological recordings were targeted to AM while animals were performing the imaging-based BMI task, with control regions in anterior motor cortex. Inset shows example waveforms from three isolated units (fast spiking unit in red). **B.** Example spiking during three successful trials (trials denoted in blue, hits denoted with pink arrows). At top, traces, from top to bottom, are: pupil diameter, running velocity, and visual cursor position. **C.** Normalised tuning to cursor positions for all single units. Each row represents the normalised firing responses to each of eight cursor positions for every recorded unit in the task (top, N = 131 units) and playback (bottom, N = 128 units) conditions. Firing responses were taken as the average firing rate for a period from 80 to 200 ms from the onset of the cursor presentation. **D.** Average population firing rates for each cursor position during task performance (red) and passive playback (blue). Shaded regions indicate 95% confidence levels. **E.** (Left) Mean firing rate for regular spiking (RS) units to different cursor positions during task performance (95% confidence interval indicated by shading, n = 7 mice). (Right) Mean firing rate for fast spiking (FS) units to different cursor positions during task performance (95% confidence interval indicated by shading, n = 7 mice).

Responses were dependent on the preceding cursor position—firing to the cursor positions closest to the target was higher if the cursor swept towards the target, but lower if the cursor swept away from it. The opposite was true for the cursor positions farther from the target: firing was higher if the cursor swept away from the target (Figure 6A-B). We surmised this might allow downstream areas to more effectively decode the cursor identity for more effective behavioural performance, and so performed a classifier analysis on the population neural responses (Meyers, 2013). Indeed, the cursor identity could be much more effectively decoded from AM neural responses during task performance vs. playback (Figure 6C). During the task, the classifier could perform above chance (12.5%) even before the cursor was present, suggesting that the neural responses also encode intention or expectation. In the passive playback condition, the classifier only performed above chance after the cursor had been presented. However, the classifier failed to decode the direction that the cursor was traveling in (towards or away from the target) in either the task or playback conditions (Figure 6D).

**Figure 6.**
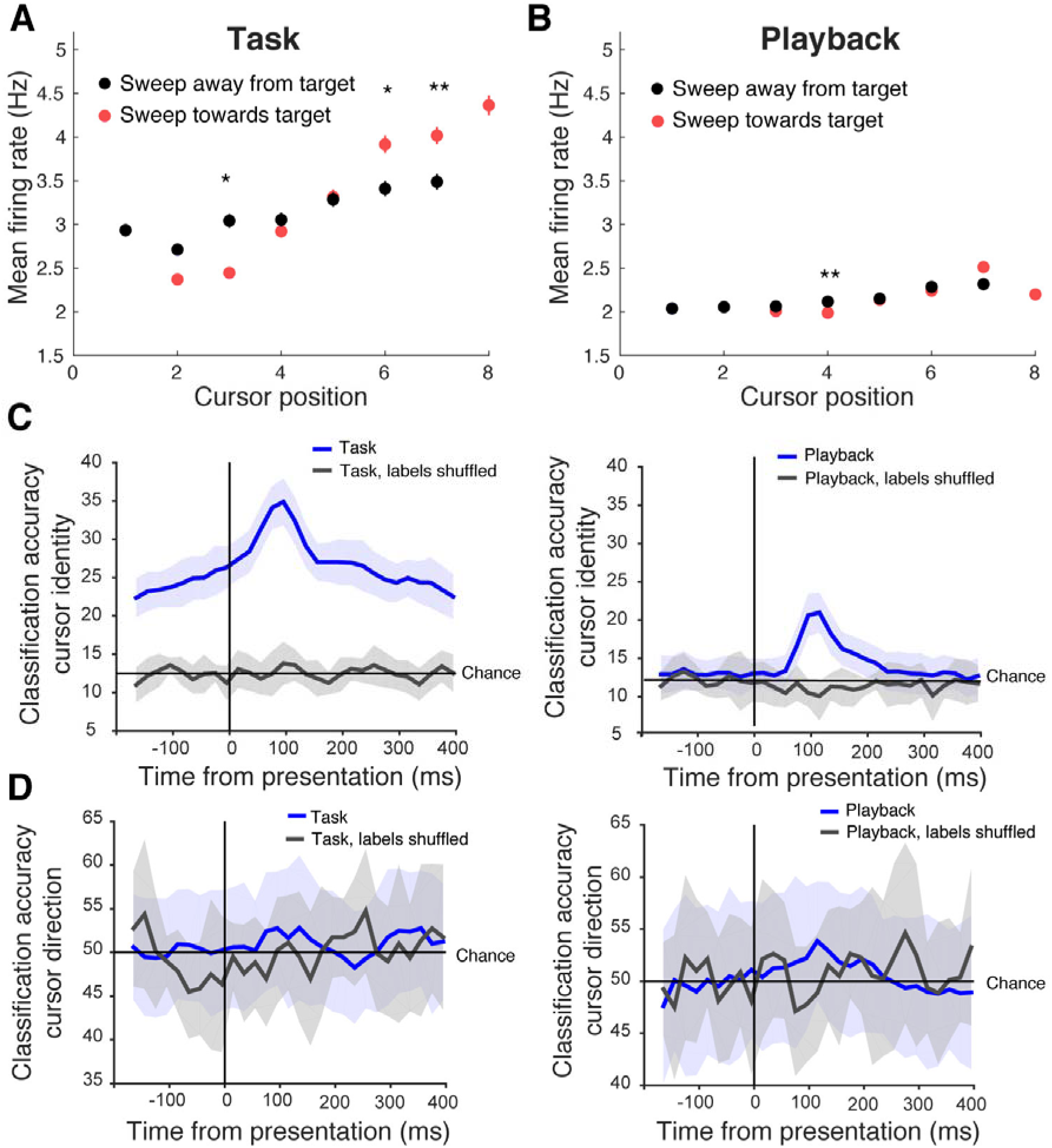
Cursor identity was more decodable from neural activity during task performance. **A.** Mean firing rate to different cursor positions depended on whether the preceding cursor position was sweeping towards (red) or away (black) from the direction of the target during task (t-test, Bonferroni corrected, * indicates p<0.005, ** indicates p<0.01). **B.** Cursor sweep direction had little effect on firing rates when animals were passively viewing playback. **C.** Classification accuracy for cursor identity, from population responses trained on real (blue) and shuffled data (grey), for neural responses during task performance (left panel) and passive playback (right panel). Shaded regions denote 95% confidence interval. During task performance, the classifier could infer the upcoming cursor position even before presentation (chance level at 12.5%), and rose higher after presentation, suggesting that neural responses encode expectation. This was not true of the passive playback condition, in which case the classifier only performed above chance during the cursor presentation period. **D.** Classification accuracy for cursor direction, from population responses trained on real (blue) and shuffled labels data (grey), for neural responses during task performance (left panel) and passive playback (right panel). Shaded regions denote 95% confidence interval. A trained classifier could not perform above chance (50%) in predicting, from firing alone, the direction that the visual cursor was moving, in either the task or playback conditions.

Pupil diameter and running velocity were significantly decorrelated during task performance compared to playback (Supp. Figure 5F), suggesting distinct mechanisms underlie pupil size modulation in the two conditions. At the population level, firing rates were uncorrelated with both pupil diameter and pupil position during the task, while firing rates were weakly correlated with pupil diameter during playback (Supp. Figure 5G-H). The fluorescence activity of the control regions was also uncorrelated with running velocity and pupil diameter (Supp. Figure 5I-J).

To understand the relationship between the firing rate of individual neurons and dorsal cortex-wide activity, we correlated the spike trains of individual units with fluorescence activity across the brain to build affiliation maps for each unit (Xiao et al., 2017; Clancy et al., 2019; Barson et al., 2018; Peters et al., 2019). We aligned these maps to the common coordinate framework of the Allen Brain Atlas using stereotaxic marks on the skull, and sorted these maps by units’ preference for different cursor locations. During task performance, the cells most responsive to the target and target-adjacent cursor position were significantly more correlated with activity across the dorsal cortex (Supp. Figure 6A). This could mean that the boosting around the target might be the result of a cortex-wide signal, or that units tuned to the target correlate more strongly with the rest of dorsal cortex during task performance, but not playback (Supp. Figure 6B-C).

## Discussion

The idea of representation is fundamental to the idea of computation, and the neocortex appears to employ hierarchies of transformed representations. It has become increasingly clear that cortical sensory representations are not strictly veridical reproductions of the outside world, but are shaped by a subject’s internal states and goals. We sought to understand how having causal control over an external object affects the cortical sensory representation of that object, given that fluent control must be informed by a constant dialogue between action and sensation (Haggard, 2017). Animals learned to causally control an external object using neural calcium signals recorded by widefield imaging, and did so by discovering and exploiting experimentally-defined mappings between their neural activity and the visual feedback that led to reward. This technique enabled us to identify cortical areas involved in task performance and then to target recordings from individual cells in these areas while animals were engaged in the task. We found that higher visual cortical areas, including area AM, were recruited during expert BMI control, and that single units in AM encoded the same visual cursor differently depending on whether the animal was causally controlling it, or passively viewing it. These results lends evidence to the idea that activity in AM, a potential homolog of parietal cortex, encodes a subject’s intention and self-monitoring of sensorimotor transformations (Andersen and Buneo, 2002; Andersen and Cui, 2009; Desmurget et al., 2009; Aflalo et al., 2015; Cui, 2016).

Previous work indicates that subjects can learn to control neuroprosthetic devices using single cells or bulk electrophysiological signals (Fetz, 1969; Bakay and Kennedy, 1998; Nicolelis, 2001; Serruya et al., 2002; Carmena et al., 2003; Weiskopf et al., 2003; Sitaram et al., 2007; Koralek et al., 2012; Hochberg et al., 2012; Collinger et al., 2013; Clancy et al., 2014; Sadtler et al., 2014; Prsa et al., 2017; Sitaram et al., 2017; Trautmann et al., 2019), but this is the first work, to our knowledge, to employ control using population calcium signals. This technique allowed us to monitor much of the dorsal cortical network as animals learned neuroprosthetic control, whereas previous BMI work has been limited to recording from neighbouring neurons (Koralek et al., 2012; Clancy et al., 2014; Sadtler et al., 2014; Prsa et al., 2017). This lends evidence to the idea that using aggregate population signals (for example, from infrared imaging (Shevelev, 1998; Abdelnour and Huppert, 2009)), rather than electrophysiological recordings from individual neurons to manipulate neuroprosthetic devices might afford more stable and minimally invasive control, robust to losing signals from, or damage to, individual control cells. We also found that the spontaneous activity correlations between the control regions, and not their spatial distance, was predictive of the animals’ fluency of performance, instructive for BMI design considerations, and similar to what has been found in other work assessing the limits of BMI learning (Sadtler et al., 2014; Clancy et al., 2014; Oby et al., 2019).

In order to learn the arbitrary action-outcome relationships required to perform BMI tasks, animals must match internally generated actions or activity with their sensory consequences. To probe how animals learned these contingencies, we changed the regions that controlled the BMI between and within training sessions: meaning that animals could not rely on a habitual activity pattern or strategy, but had to continually explore different neural patterns to achieve reward on different training days. Animals did so by ‘exploring’ with highly variable neural activity patterns early in a training session, until they discovered a successful activity pattern they could reliably exploit (Figure 3). Target hit rates dropped when reward was dispensed randomly, unlinked to the target zone, indicating the animals’ task performance was goal-directed, and not habitual.

Animals could decorrelate normally correlated brain areas during task execution, even though spontaneous activity was widely correlated across cortex in mice not performing the task, in agreement with previous work (Ledochowitsch et al., 2013). We found this to be true of both anterior and posterior cortical areas—in mice trained to control an auditory cursor, for example, posterior visual control areas could also be decorrelated during the task (Figure 3B). However, animals performed better when control regions were less spontaneously correlated (Figure 3C). Pupil and locomotion also became significantly decorrelated during task performance, indicating that task-engagement and locomotion may engage distinct arousal mechanisms (Supp. Figure 5, Vinck et al., 2015; Reimer et al., 2016; Clancy et al., 2019).

By imaging dorsal cortex as animals performed this task, we were able to screen for cortical areas engaged during expert BMI control. On the first day of training, V1 was most active during the task, but as animals learned the task over days, higher visual areas, including AM, PM and RL, became more active as animals controlled the cursor. When the same cursors were played back to animals in an open-loop fashion (e.g. not controlled by animals), activation was again mainly evident in V1, suggesting these higher areas were involved in the goal-directed aspect of task performance. AM and RL are considered by some to be rodent homologues of parietal cortex (Glickfeld and Olsen, 2017; Wang et al., 2011), which has been shown in humans to be involved in intention and monitoring the mapping between action and outcome (Andersen and Buneo, 2002; Desmurget et al., 2009; Andersen and Cui, 2009; Aflalo et al., 2015; Cui, 2016). However, it was unclear whether the recruitment of these higher visual areas over learning was related to the fact that they are involved with sensorimotor transformations generally, or because they’re involved more specifically in planning and intentional control.

In order to clarify the role of these higher visual areas in causal control, we targeted extracellular recordings in one of the functionally identified task-active areas, AM, to probe task-related changes in the spiking of single units. We found that units were more active during task performance than during passive cursor playback of the animal’s recent performance. In particular, units were more responsive to the cursor when it was at the target, and target-adjacent, positions during the BMI task compared to passive playback, similar to spatial attention boosting evident in previous work (Moran and Desimone, 1985; Engel et al., 2016), and in accordance with evidence that attention can reshape stimulus representations in a manner that more effectively guides decisions (Ruff and Cohen, 2019). Indeed, we found that a classifier trained on neural responses to the different cursor positions could effectively classify responses to these visual cursors, and could do so with significantly better accuracy when the animal was actively engaged in the task versus passively viewing a playback of the same cursors. In the task condition, the classifier ramped up in accuracy even before the cursor was presented, suggesting that neural responses were prospective, but whether this represents an intention (e.g. a control signal) or an expectation (e.g. a passive prediction) remains to be determined in future work. This boosting did not reflect reward expectation, and the animal did not use saccades or overt movements to perform the task. This boosting was also sensitive to the task goal: if the cursor was positioned close to the target and sweeping towards it, responses were boosted relative to when it was sweeping away. If the cursor was far from the target and sweeping farther away from it, responses were also boosted relative to the cursor at the same position sweeping towards the target. This suggests that firing rates reflect intention, and may also reflect a valence of the animal’s perceived fluency of cursor control—that is, whether it was successfully or unsuccessfully guiding the cursor towards its intended goal (Lee and Dan, 2012).

We present a novel task and imaging method for exploring the encoding of action-outcome assessments, which allowed us to simultaneously monitor–with both dorsal cortex-wide and cell-level resolution–what activity patterns support the causal control of external objects. We believe imaging-based BMIs can serve as a useful paradigm for studying the cortical dynamics involved in flexible learning, affording the experimenter control of what is learned, and by what areas or cell types. This makes it an excellent system for studying how the brain deals with sensory and rule manipulations, as well as a promising platform for studying credit assignment. It’s also a promising proof-of-principle for exploring the potential–and limitations–of future imaging- and population-activity-based BMIs. While using widefield imaging afforded us a view of the dorsal cortex as animals learned neuroprosthetic control, there are a number of limitations to this method. While widefield fluorescence signals largely reflects neural firing, there is also a contribution from neuropil, as well as hemodynamic effects that were uncorrected in this study, as in our previous work we found it did not significantly impact our findings (Clancy et al., 2019). However, there is clearly some component of the signal contributed by blood vessels (e.g. in Figure 3B), so this is a limitation of the study.

Another limitation is that widefield imaging is limited to recording from dorsal cortex. Cortical neuroprosthetic control requires interactions with basal ganglia (Koralek et al., 2012; Neely et al., 2018), from which we cannot record using this method. Furthermore, we know from work in humans that prefrontal cortex (PFC) is involved in the sense of control over external stimuli, but we cannot record signals from PFC using this preparation due to its obscuration by the frontal sinus.

While we found increased evoked spiking to a visual cursor in the target location using this preparation, we do not know the exact cellular or neuromodulatory mechanisms giving rise to this difference. We found task-related increases in activity emerge over learning specifically in AM, PM and RL, and are not present during passive visual playback or when visual feedback was randomised. However, as we only recorded neuronal responses in AM, we do not know if the changes in neural tuning in AM are specific to this area, nor whether they are specifically required for task performance. Future work may address this by using mesoscale 2-photon imaging to record from or manipulate activity in molecularly defined neural subpopulations in parietal and control areas during task performance.

## Acknowledgements

The authors would like to thank Adrianne Zhong for help with the camera control code, Ivana Orsolic for help with the widefield imaging setup, Lisa Hoermann for performing surgeries for this project, and Sonja Hofer, Petr Znamenskiy, Spencer Wilson, Alex Naka, and Adrei Khilkevich for manuscript feedback and discussions of this work. This work was supported by the European Research Council (NeuroV1sion 616509 to T.D.M-F.), Swiss National Science Foundation (SNSF 31003A_169802 to T.D.M.-F.), the Wellcome Trust (090843/E/09/Z core grant to the Sainsbury Wellcome Centre), the EMBO Long-term Fellowship (ALTF 1481-2014 to K.B.C.), the HFSP Postdoctoral Fellowship (LT000414/2015-L to K.B.C.), and the Branco Weiss-Society in Science grant (K.B.C.).

## Author contributions

K.B.C. planned and executed the experiments and analysed the data. K.B.C. and T.D.M.-F. wrote the manuscript.

## STAR Methods

### Resource availability

#### Lead contact

Further information and requests for resources and reagents should be directed to Kelly Clancy, the lead contact.

### Materials availability

This study did not generate new unique reagents or mouse lines.

### Data and code availability

The data and code used in this study are available from the lead contact upon reasonable request.

### Experimental model and subject details

#### Mice

All experimental procedures were carried out in accordance with institutional animal welfare guidelines and licensed by the Swiss cantonal veterinary office. TRE-Gcamp6s mice (Wekselblatt et al., 2016) (Jackson laboratories, https://www.jax.org/strain/024742) were crossed with B6.CBA-Tg(Camk2a-tTA)1Mmay/DboJ mice (Jackson laboratories, JAX 007004), to drive the expression of gCamp6s in CamKII+ pyramidal neurons. Animals were housed in a facility using a reversed light cycle, and recordings were taken during their active period. Eleven female mice were trained on the task, and we took electrophysiological recordings from seven of these, ranging between P55-P75. All animals were healthy, had never undergone previous procedures, and ranged in weight from 15-19g. Sample sizes were not statistically determined, but were consistent with previous papers using related methodology (Clancy et al., 2019; Xiao et al., 2017). Animals were group housed with same-sex littermates in enriched environments (including running wheels, cardboard tubes and chewing toys).

### Method details

#### Surgery

A week before training, mice were prepared for imaging. Animals were anaesthetised with a mixture of fentanyl (0.05 mg per kg), midazolam (5.0 mg per kg), and medetomidine (0.5 mg per kg). The animal’s scalp was resected and a head plate was secured to the skull. Four stereotaxically placed marks were made to enable alignment of the imaged brain with the Allen Brain Atlas (http://mouse.brain-map.org/static/atlas) post hoc, using the Allen Brain API (brain-map.org/api/index.html). The exposed skull was cleaned and covered with transparent dental cement to avoid infection, and to cover the cut scalp edges (C&B Metabond). This was polished to enhance the transparency of the preparation. A custom-made 3D printed light shield was cemented to the skull and head plate to avoid light leaks from the visual feedback presented on two computer monitors.

#### Behavioral setup and recordings

The recording chamber was sound-isolated and shielded from outside light. Mice were head-fixed under the microscope and free to run on a Styrofoam running wheel (diameter = 20 cm, width = 12 cm). The animals’ movements were recorded using a rotary encoder in the wheel axis (pulse rate 1000, Kubler). Two monitors were placed side by side in front of the mouse, angled towards one another (21.5” monitors, ~20 cm from mouse, covering ~100×70 degrees of visual space), similarly to the setup described in (Poort et al., 2015). A reward port was place in front of the animal, where reward delivery was triggered via pinch solenoid one second after target hit (NResearch) and animal licks were detected using a custom piezo element coupled to the spout. All behavioral data were recorded using custom MATLAB software and a PCI-6320 acquisition board (National Instruments).

On electrophysiological recording days, pupil recordings were taken by illuminating the animal’s right eye with a custom IR-light source and imaging with a CMOS camera (DMK22BUC03, Imaging Source, 30 Hz) using custom MATLAB software. Pupil size was determined as described in (Orsolic et al., 2019): images were first smoothed with a 2-D gaussian filter and thresholded to low luminance areas. These thresholded regions were then filtered by circularity and size to automatically detect the pupil region. Pupil edges were detected using the canny method, and ellipses were iteratively fit to the region, tasked to minimise the geometric distance between the area outline and the fit ellipse using nonlinear least squares (MATLAB function fitellipse, Richard Brown). The pupil diameter was taken to be the major axis of the ellipse, then normalised by animal. Pupil recordings from one animal had to be discarded, as the video was not sufficiently in focus.

#### Behavioral training

After recovery, mice were acclimatised to head fixation for a minimum of two days, and started on food restriction. Awake animals were head-fixed under the microscope and free to run on a Styrofoam wheel. A baseline of spontaneous activity was taken on every training day (10-20 minutes) in order to estimate spontaneous hit rates. The decoder was calibrated such that animals achieved ~25% performance on their first day. Two small control regions were chosen for real-time read out. In the case of visual feedback task, these were all located in primary and secondary motor cortex, avoiding ALM. In the auditory feedback task, control regions were placed in posterior cortex, over visual and retrosplenial areas. The placement of the two control regions was usually ipsilateral but sometimes contralateral to each other. The same control regions were used for the first few days of training, then changed from day to day, or within sessions, so that animals did not learn a fixed control strategy (see Table 1).

Activity was imaged at 40 Hz and the mean fluorescence from each control region was transmuted to the cursor’s position on screen with a simple transform:

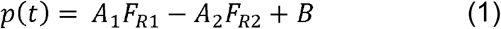

where p is the cursor position, F_R1_ and F_R2_ are the instantaneous fluorescence (ΔF/F) of control regions one and two, respectively, and A_1_, A_2_ and B are coefficients set based on the daily spontaneous baseline recordings (minimum 10 minutes). P was rounded to the nearest integer to determine the discrete cursor location. A1 and A2 were determined by dividing the full dynamic range of each recorded area during the baseline by half the number of cursor positions:

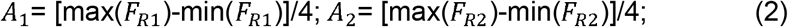

B represents the activity baseline of both areas. The chance performance was then assessed by running the baseline data through the decoder to estimate how often the animal would have achieved the target with spontaneous activity.

ΔF/F was calculated using a moving baseline, set as the tenth percentile of points from the preceding 20 seconds of data. The raw fluorescence was converted to ΔF/F using a moving baseline of 5 minutes of activity. The display updated at approximated 10 Hz, with a latency of 300 ms from camera to screen, measured using a photodiode placed on one of the monitors (Thorlabs, PDA100A-EC). Activity in R1 would cause the cursor to move towards the target location in the centre of the animal’s visual field, while increased activity in R2 would cause the cursor to move away from the target zone. The cursor was presented on two monitors so that the animal could track the cursor with both eyes; its goal was to move the cursors presented on the two screens on either side to the middle of its visual field. These changes were binned, such that the cursor could take one of eight possible locations on the screen. The cursor had to be held at the target position for 0.3 seconds to count as a hit, at which point the cursor disappeared. When a target was hit, a MATLAB-controlled Data Acquisition board (National Instruments, Austin, TX) triggered the administration of a soyamilk reward following a 1 second delay. The next trial could be initiated within 5 seconds of reward delivery, but only when the activation of R1 relative to R2 returned to the mean value recording during spontaneous activity (to ensure enough time had passed for large transients to decay, given slow calcium dynamics). This was return to baseline condition was rarely triggered (~5% of trials) and on average lasted under 2 seconds. If the animal did not bring the cursor to the target within a 30 second trial, the cursor disappeared, and the animal received a white noise tone and a 10 second ‘time out.’

We trained a separate cohort of four mice using an auditory, rather than visual, feedback cursor, where activity was transmuted to the pitch of a feedback tone (Clancy et al., 2014). As with the visual feedback task, a spontaneous baseline was recorded every day (10-20 minutes) to assess chance levels of performance and calibrate the decoder. Activity from two arbitrarily chosen regions was entered into an online transform algorithm that related neural activity to the pitch of an auditory cursor:

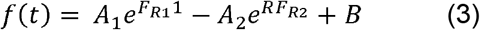

Where f is the cursor frequency, F_R1_ is the instantaneous ΔF/F of R1, F_R2_ the instantaneous ΔF/F of R2, and A1, A2, and B are coefficients set based on the daily baseline recording. As above, ΔF/F was calculated using a moving baseline, set as the tenth percentile of points from the preceding 20 seconds of data. Linear changes in firing rate resulted in exponential changes in cursor frequency, and frequency changes were binned in quarter-octave intervals to match rodent psychophysical discrimination thresholds. As with the visual task, a trial was marked incorrect if the target pitch was not achieved within 30 seconds of trial initiation. The auditory feedback was played using speakers mounted on 2 sides of the imaging platform.

#### Task and control conditions

In the **task condition**, the position of the presented cursor was determined by the control regions’ instantaneous activity, according to equation (1) above. Rewards were given when the cursor hit the target zone (cursor position 8). Similarly, in the **random reward condition**, intended to test whether animal’s task engagement was goal-directed or habitual, the position of the presented cursor was determined by the control regions’ instantaneous activity, according to equation (1) above. However, here rewards were not linked to the cursor position, but given out at random time intervals at a rate matched to an expertly performing animal (approximately 1.5 rewards/minute). In the **random feedback condition**, intended to confirm that the animal was using the visual feedback to inform its behaviour, the position of the presented cursor was not linked to the control regions’ instantaneous activity, but instead was drawn randomly from a Gaussian distribution matching the mean and variance of a typical task condition. In the **passive playback condition**, the presented cursor position was no longer linked to the control regions’ activity, but was purely a replay of the cursor positions from the training session the animal had undergone previously. Thus, the cursor positions and timing of cursor presentations were matched to those of the task condition.

#### Widefield imaging

Widefield imaging was performed through the intact skull using a custom-built epifluorescence macroscope with photographic lenses in a face-to-face configuration (85mm f/1.8D objective, 50mm f/1.4D tube lens; (Ratzlaff and Grinvald, 1991)). Data were recorded using a CMOS camera (Pco.edge 5.5, PCO, Germany) in global shutter mode. 16-bit images were acquired at a rate of 40 Hz and binned 2×2 online using custom-made LABVIEW software. A constant illumination at 470 nm was provided (M470L3, Thorlands, excitation filter FF02-447/60-25), with average power ~0.05 mW.mm2 (emission filter 525/50-25, Semrock). The imaging site was shielded from light contamination using a 3D-printed blackout barrier glued to the animal’s skull. Signals from the two control regions were sent via UDP to a computer providing visual or auditory feedback to the mouse, using custom MATLAB software.

#### Electrophysiological recordings

The day before recording, mice were anesthetised with isofluorane and a small craniotomy was opened over AM, which was functionally identified during task performance, and stereotaxically confirmed. The craniotomy was kept damp with Ringer’s solution and sealed with KwikSil (World Precision Instruments). Recordings were taken on the following day to avoid residual effects of anesthesia.

On the recording day, animals were head-fixed under a custom-built widefield microscope, the skull and cortex were cleaned with Ringer’s solution, and the KwikSil plug removed from the craniotomy. A custom-designed silicon probe (64 channels, 2 shanks, Neuronexus, as described in (Clancy et al., 2019)) was inserted at an angle of ~45 degrees from normal to cortex. The probe consisted of two shanks with 64 sites total, organised into 16 ‘tetrodes’, each consisting of 4 sites located 25 um apart from each other within-tetrode, and tetrodes spaced 130 um apart from each other. A small amount of KwikSil or agar was used to cover the exposed cortex after the probe was in place. After allowing the probe to settle for 20-30 minutes, neural activity was recorded using the OpenEphys recording system (Siegle et al., 2017). Behavioral and stimulation data, including pulses representing each camera frame, were recorded using OpenEphys, enabling the alignment of electrophysiological signals with imaging and behavioral data. Ephys recordings were filtered between 700 and 7000 Hz, and spikes detected using the Klustakwik suite (Schmitzer-Torbert et al., 2005). Clusters were assigned to individual units by manual inspection, excluding any units without a clear refractory period. Units were separated into fast and broad spiking units by their peak-to-trough time, using a cutoff of 0.66 ms (Barthó et al., 2004).

#### Data analysis

Raw imaging data were checked for dropped frames, spatially binned 2×2, and loaded into MATLAB as a mapped tensor (Muir and Kampa, 2015). The raw fluorescence was converted to ΔF/F using a moving baseline, calculated as the tenth percentile of points from the preceding 20 seconds of data. We did not perform hemodynamic correction as previous work indicates that hemodynamic and flavoprotein signals contribute minimally compared to the calcium responses (Vanni and Murphy, 2014; Xiao et al., 2017; Clancy et al., 2019).

Task-activation maps were calculated by taking the normalised average of fluorescence movies during the task, or visual cursor playback period, subtracted by periods when animals were not performing the task or viewing any visual stimuli (including periods of spontaneous activity, and reward collection). To ensure that differences between early and late in training were not influenced by possible differences in the statistics of the visual feedback cursor, we randomly excluded success trials on late training days in order to have comparable numbers of success and failure trials between early and late training, however, including or excluding these trials did not influence the result. To build the single-unit affiliation maps (Supp. Figure 6, see also (Clancy et al., 2019)), spike trains were binned to match imaging frames, and maps were calculated by taking the correlation of each unit’s spike train with each pixel’s ΔF/F.

Spectral entropy was calculated in 10 second windows, each overlapping by 5 seconds. The calcium signal of the control areas was transformed into power spectral density (PSD) during these windows (the magnitude squared of the signal’s Fourier transform). This was then used to calculate the spectral entropy for that time span:

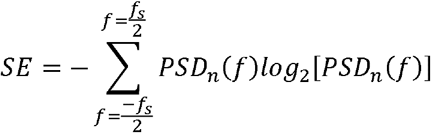

Where SE is the spectral entropy, PSD_n_ is the normalised PSD, and *f* is frequency.

#### Classifier analysis

Spike data were binned into 50 ms bins, and split into 40 segments for training: 39 of these splits were used for training the classifier and 1 was used for testing. The data were z-score normalized so that high firing rate units didn’t bias the classifier results. This data was then used, along with the cursor position labels, to train the classifier: a mean vector was created for each cursor position class based on the training data, and predictions were made on the test data by choosing the label class with the maximum correlation between the test and training mean vectors. These predicted labels were compared with the true labels to generate an average classifier accuracy over each tested time bin. This process was repeated 20 times using different training/test splits to cross-validate the results. The final reported classification accuracy is the mean of these 20 runs.

### Quantification and statistical analysis

All data were analysed using custom code in MATLAB. The statistical tests used in our analyses are indicated in the figure legends, which also includes the value of n, and whether it refers to animals or single units. Differences were tested using Student’s t-test, and Bonferroni corrected where apropriate. P-values are reported in the figures as well as legends, and significance is herein defined as p less than 0.05. The pupil video for one animal, taken on the final day of recording, had to be excluded as it was too out of focus to determine the pupil size. Sample sizes were not statistically determined, but were consistent with previous papers using related methodology (Clancy et al., 2019; Xiao et al., 2017). Data were assumed to be normal but this was not systematically tested. Animals were trained in three separate cohorts (the first, a cohort of four animals trained on the auditory feedback task, the second, a cohort of 5 animals trained on the visual feedback task, and the third, a cohort of 2 animals trained on the visual feedback test) as an internal replication.

## Supplemental Information

The sensory representation of causally controlled objects

**Supplemental Figure 1.**
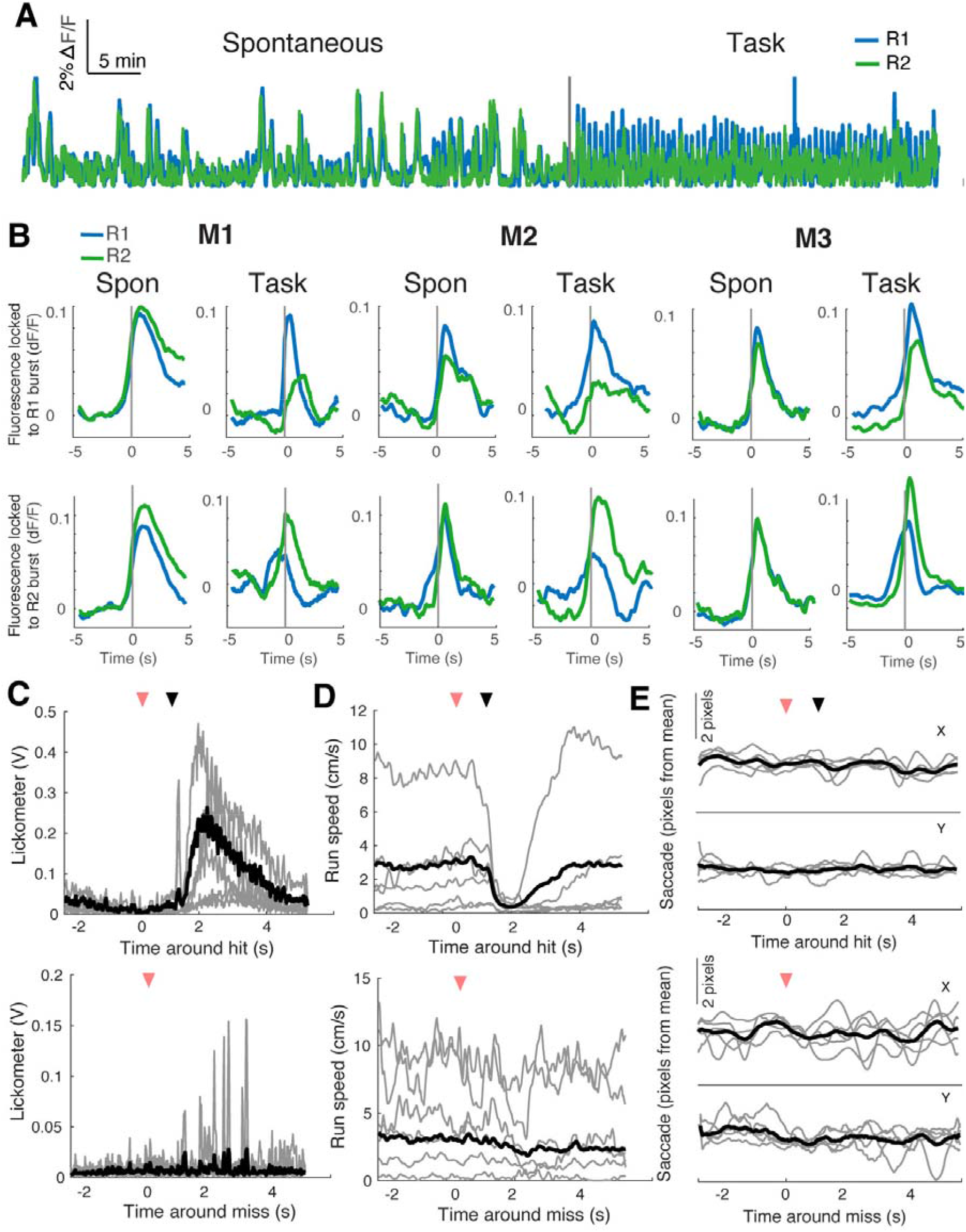
Animals did not use gross motor movements to perform task (related to Figure 1) **A.** Example fluorescence trace from R1 and R2 in pre-task spontaneous activity, and during task, indicating a clear change from spontaneous patterns. **B.** Analysis of activity throughout the task indicated that activity in the two control regions generally became decoupled during task performance, and not just around hits. Average ΔF/F in the control regions triggered by bursts in R1 (top row) or R2 (bottom row), for the same three example animals, for spontaneous activity (left) and during the training session (right). The burst threshold was taken as the z-scored value of ΔF/F > 3, in order to capture large events. **C**. Lick averaged around hits (top panel) and misses (bottom panel) on a day in late training (n = 7 mice, grey traces indicate individual mice, black trace indicates mean). Pink arrow indicates target hit (top panel) or time out (bottom panel), black arrow indicates reward delivery (top panel). **D.** Animal’s running speed, averaged around hits (top panel) and misses (bottom panel) on a day in late training (n = 7 mice, grey traces indicate individual mice, black trace indicates mean). Pink arrow indicates target hit (top panel) or time out (bottom panel), black arrow indicates reward delivery (top panel). **E**. Eye saccades averaged around hits (top panel) and misses (bottom panel) on final day of training (n = 6 mice). Average movement is shown for both x and y directions of pupil image. Pink arrow indicates target hit (top panel) or time out (bottom panel), black arrow indicates reward delivery (top panel).

**Supplemental Figure 2:**
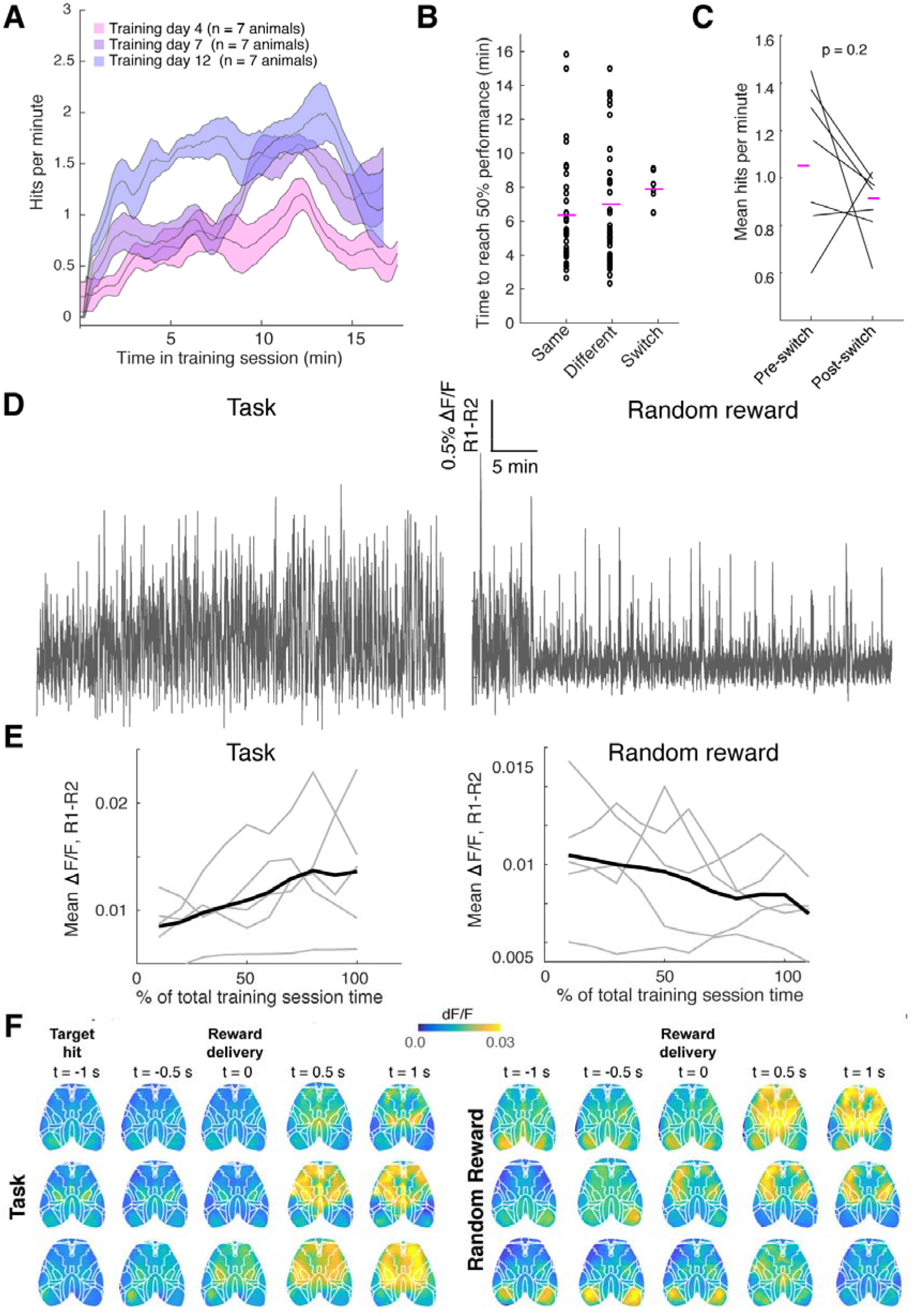
Animals exhibit quasi-intentional, specific neural activity control during task performance (related to Figure 2) **A.** Hit rate over the course of a training session averaged over 7 mice for three training days (shaded area indicates s.e.m., 7 animals for each day). **B.** Once animals were performing above chance (from day 4 on), there was no significant difference in the time it took animals to reach 50% performance whether they were using the same ROIs as the previous day of training, or different ones (t-test, p = 0.4). On day 6, when ROIs were switched mid-session, there was also no significant difference in the time it took, post-switch, for the animal to recover their performance to 50% compared to animals using the same or different control regions to the day before (t-test, p=0.3, 0.7 respectively, each dot indicates performance on one training day). **C.** On the day control regions were switched mid-session, there was no significant difference in performance between animals before their control regions were switched, and their recovery after the switch. Each line represents one mouse (n = 7 mice). **D.** Example trace of the difference between control areas’ activity during task (R1-R2) for one mouse, indicating an increase in differential modulation over the course of a training session (left). These modulations decreased when reward was provided randomly (right). **E.** The slope of the difference in the two control regions fluorescence over training was positive during the normal task condition (left, p < 0.01, light-weighted lines each represent one of n = 5 mice that underwent the randomized reward condition on day 8 of training, heavy line represents mean) and negative over the course of the randomised reward condition (right, p < 0.01, light-weighted lines each represent one of n = 5 mice that underwent the randomized reward condition on day 8 of training, heavy line represents mean). **F**. (Left) Example activity maps locked around reward delivery for three animals performing the task. One second before the reward delivery marks the target hit, after which the cursor disappeared during the 1 second wait time. After reward delivery, strong activity is evident in frontal areas when animals are licking and touching the spout. (Right) Example activity maps locked around reward delivery for three animals in the random reward condition. Rewards were given randomly, unlinked to target hits, so the visual cursor was usually present throughout the reward collection period. After reward delivery, strong activity is evident in frontal areas.

**Supplemental Figure 3.**
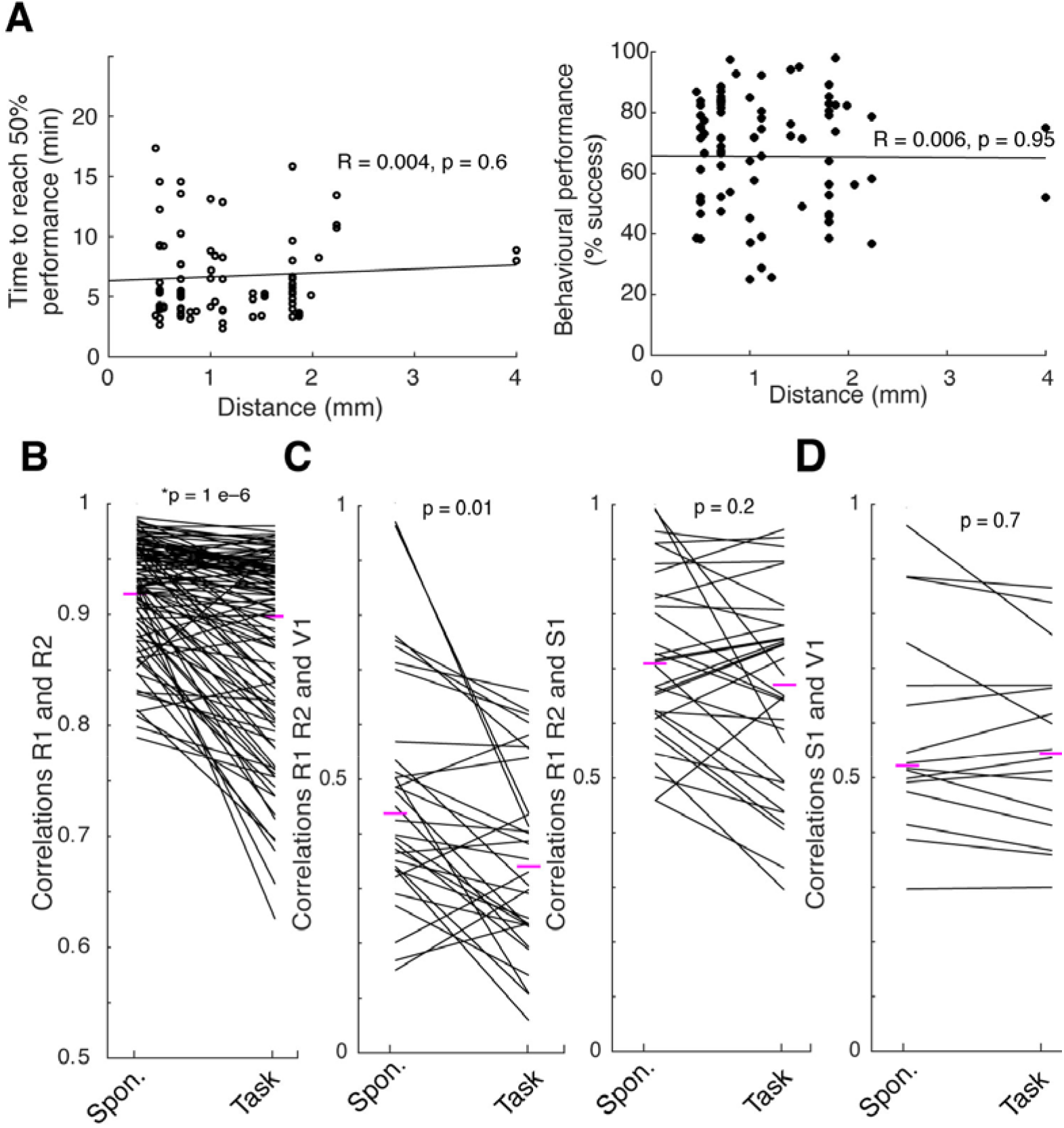
Animals could modestly decorrelate areal brain activity during task performance (related to Figure 3) **A.** (Left panel) The distance between control regions did not have an effect on the time it took animals to achieve 50% performance (for 83 training sessions, including all days of above chance performance from all 7 animals, e.g. from day 4 on). (Right panel) The distance between control regions did not have an effect on final performance (for 83 training sessions, including all days of above chance performance from all 7 animals, e.g. from day 4 on). **B.** Correlations between R1 and R2 dropped modestly from spontaneous levels during task performance (104 control region pairs from 7 animals, on all training days: on average 15 days of training per mouse, paired t-test, p = 1e-6). **C.** Left panel: correlations between control regions and activity in V1 dropped between spontaneous activity and task performance, but correlations between control regions and primary somatosensory cortex (S1) were not significantly different between conditions. Areas were identified by registering imaged brain surface with Allen Brain Atlas by stereotaxic marks; 7 animals, for expert level performance on days 9 and 14 of training paired t-test, p = 0.01, p=0.1, respectively for V1, S1). **D**. Correlations between primary visual cortex and primary somatosensory cortex did not change between spontaneous activity and task performance (areas as identified by registering imaged brain surface with Allen Brain Atlas by stereotaxic marks; 7 animals, for expert level performance on days 9 and 14 of training paired t-test, p = 0.02).

**Supplemental Figure 4.**
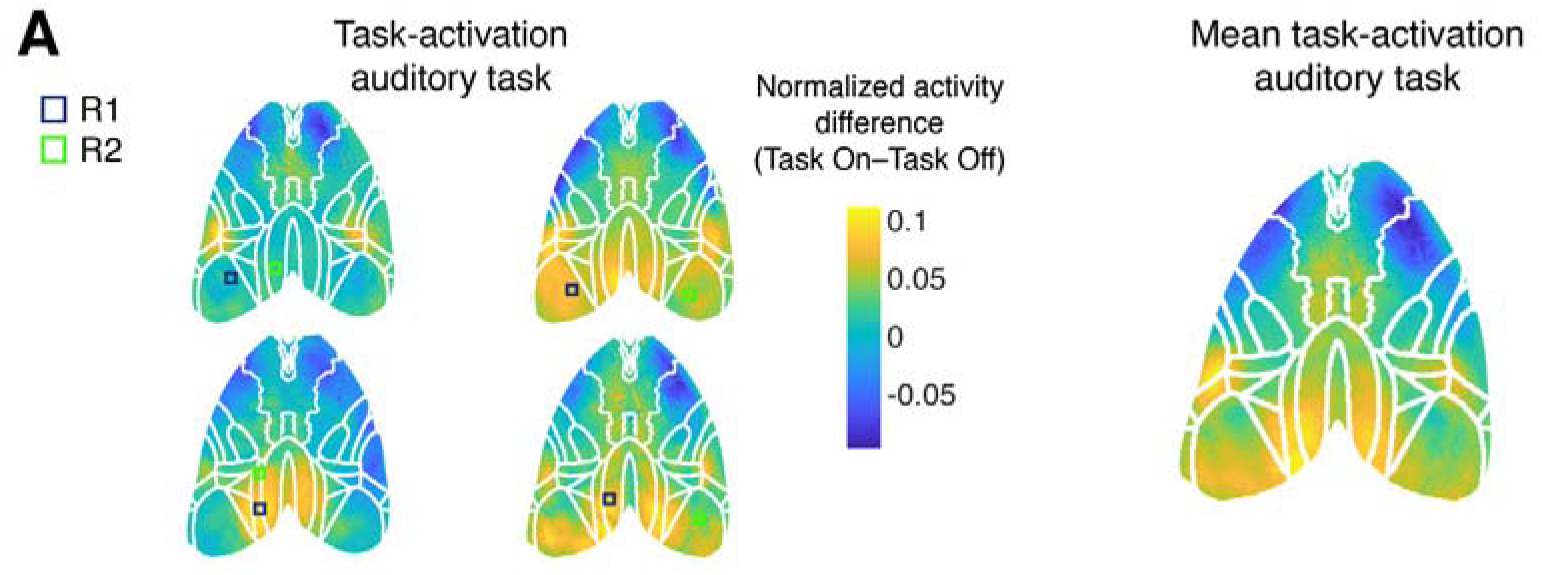
Expert animals had increased activity in higher visual areas during task performance (related to Figure 4) **A.** Left panel: example task-related activity maps from four animals trained on an auditory version of the task. Right panel: mean task-related activity map indicating involvement of RL, a lateral homolog of parietal cortex in mice and a multimodal associative area (mean from n = 4 mice). Visual areas appear task-activated (despite there being no visual feedback in this task) likely due to the fact that the control regions were placed over visual areas. Control regions are shown as slightly larger than they actually were for better visibility.

**Supplemental Figure 5.**
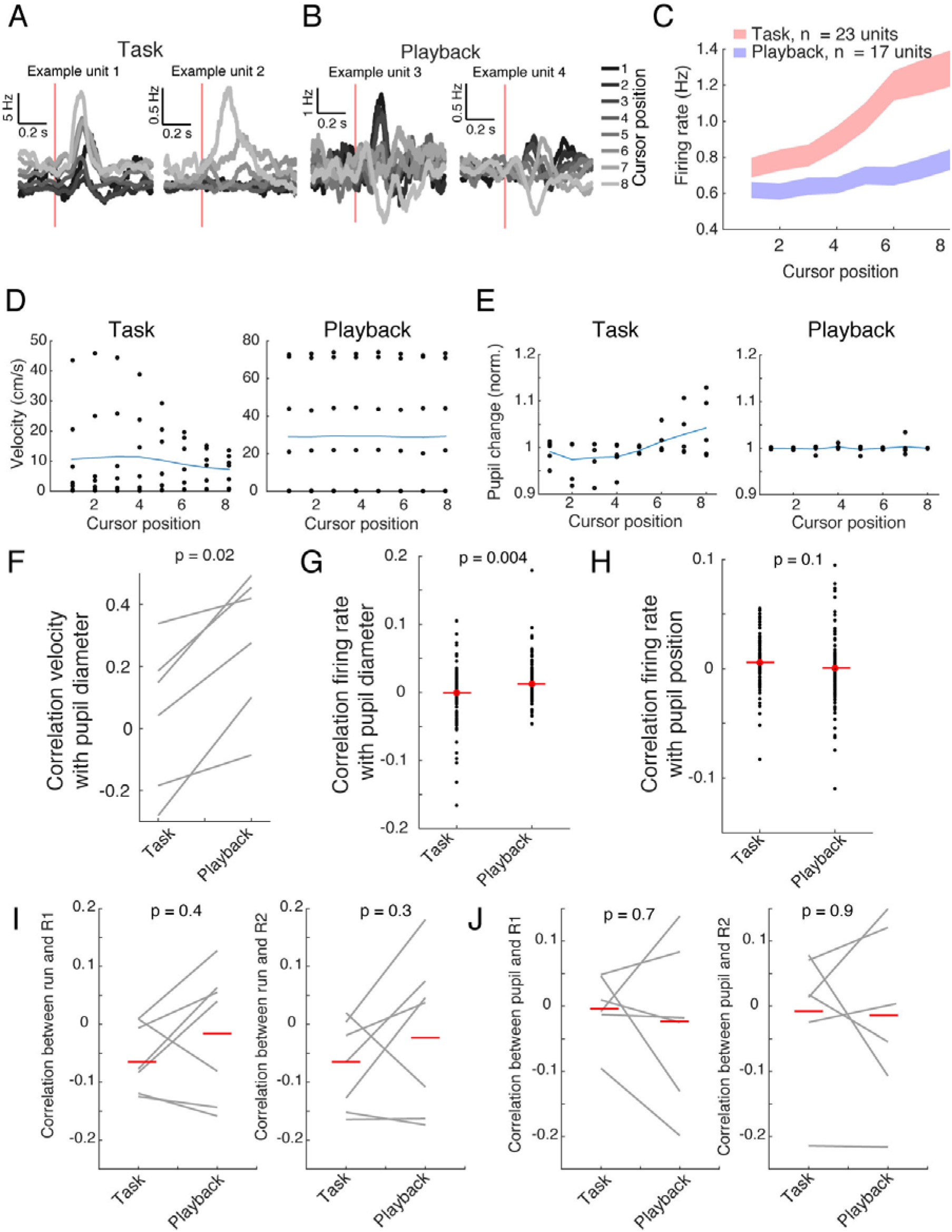
Pupil and locomotion become uncoupled during task (related to Figure 5) **A.** Responses to the visual cursor at the 8 monitor positions for two single units recorded during task performance. The red line denotes the time the cursor appeared; the trace colour denotes cursor position. Here the spiking responses have been smoothed by convolving with a gaussian kernel with standard deviation of 25 ms. **B.** Responses to the visual cursor at the 8 monitor positions for two single units recorded during passive playback. The red line denotes the time the cursor appeared; the trace colour denotes cursor position. Here the spiking responses have been smoothed by convolving with a gaussian kernel with standard deviation of 25 ms. **C.** Firing rates for units recorded in 2 animals where the target cursor position was rewarded during passive playback (blue trace). The enhanced firing for the target-adjacent cursor positions during task performance (red trace) remained, suggesting the boosting was not simply reward expectation (95% confidence interval indicated, n = 2 mice, 23 units in the task condition and 17 units in the passive playback condition). **D.** Average running velocity at each cursor location during recordings during task (left) and playback (right), (n = 7 mice, final day of recording). **E.** Normalized pupil diameter at each cursor location during task (left) and playback (right), (n = 6 mice, final day of recording). **F.** Pupil diameter and running speed were significantly decorrelated during task performance compared to playback (reward collection periods and inter-trial intervals excluded, n = 6 mice, final day of recording, each line represents correlations for one mouse.) **G.** The correlation between single unit firing rate and pupil diameter was not different from zero during task, but slightly higher than zero during playback (reward collection periods and inter-trial intervals excluded, n = 6 mice, N = 105 units task, n= 122 units playback). **H.** The correlation between single unit firing rate and pupil position was not different from zero during task or playback conditions (reward collection periods and inter-trial intervals excluded, n = 6 mice, N = 105 units task, n= 122 units playback). **I.** R1 (left panel) and R2 (right panel) activity were not significantly correlated with running velocity, either during task performance or passive playback (reward collection periods and inter-trial intervals excluded, N = 7 mice, each line represents correlations for one mouse). The correlation between R1 or R2 and run velocity during task and passive playback were not significantly different from zero (t-test: for R1, p = 0.08, p=0.6, respectively for task and playback; for R2: p = 0.08, 0.7 respectively for task and playback), nor significantly different from each other (paired t-test, p = 0.4). **J.** R1 (left panel) and R2 (right panel) activity were not significantly correlated with pupil diameter, either during task performance or passive playback (reward collection periods and inter-trial intervals excluded, N = 6 mice, t-test; each line represents correlations for one mouse). The correlation between R1 or R2 and pupil diameter during task and passive playback were not significantly different than zero (t-test, for R1: p = 0.9, 0.7 for task, playback, respectively; for R2: p = 0.9, 0.8 for task, and playback, respectively), nor significantly different from each other (paired t-test, p = 0.4).

**Supplemental Figure 6.**
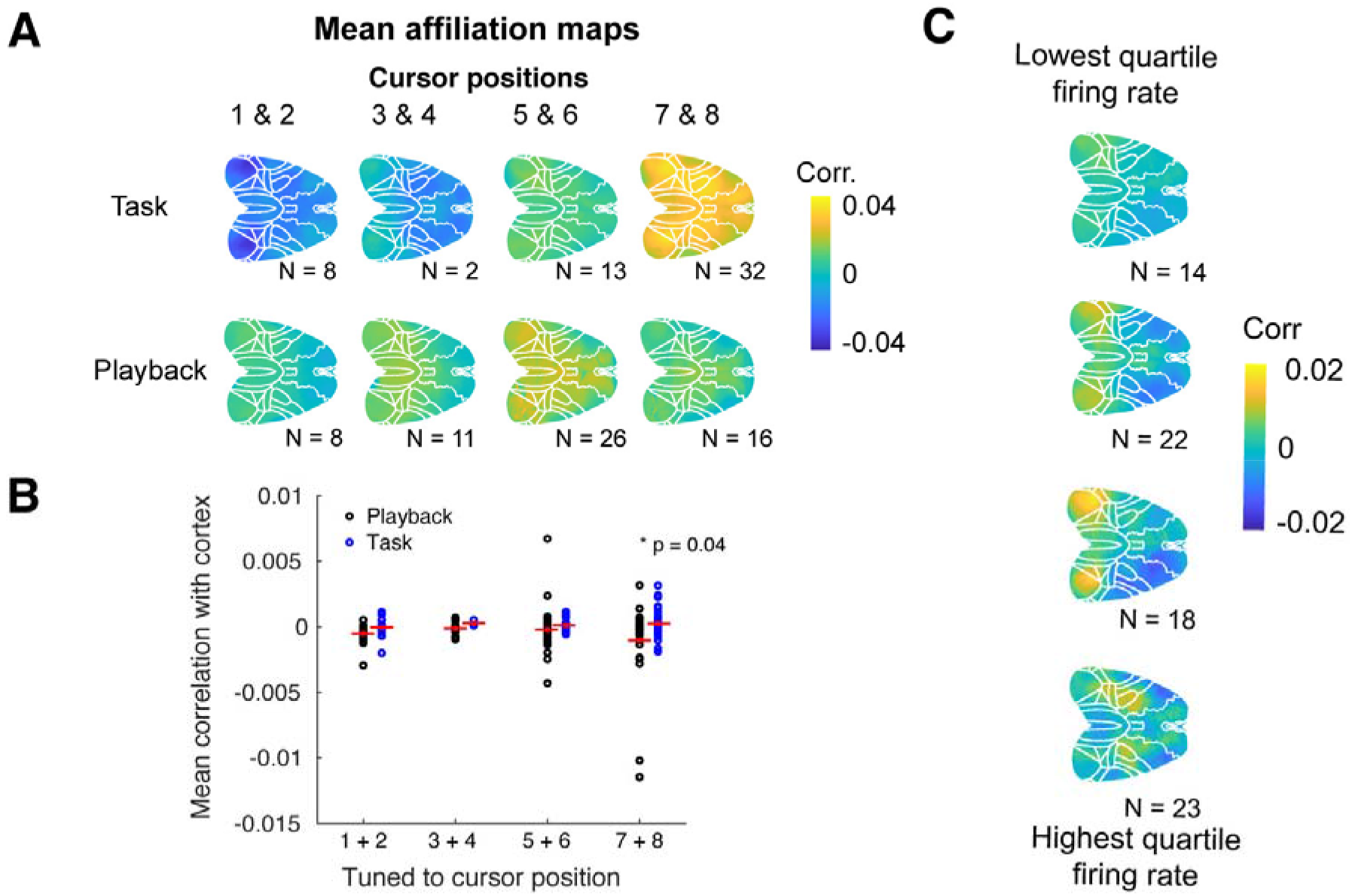
Cortex-wide affiliations of units tuned to different cursor positions (related to Figures 5 & 6) **A.** Spike trains for each unit were correlated with the activity of each pixel to build cortex-wide affiliation maps. These were sorted into bins based on which cursor position units were most responsive to. The average of these maps is shown in the top row for task performance, and the bottom row for passive playback. The reported N under each average map indicates the number of single units in that bin for that condition. **B**. Mean correlation of spiking activity with calcium activity across dorsal cortex, sorted by units’ cursor preference for task and playback. **C**. The trend evident in **A** is not due to higher firing rates of target cursor-tuned units, when units were organized into quartiles of mean firing rates rather than cursor preference. The reported N under each average map indicates the number of single units in that bin for that condition.

**Supplemental Table 1.** Control region coordinates (related to Figure 1) Related to Figure 1B: approximate stereotaxic coordinates, in millimeters, of control regions on all days of training, reported in [medial/lateral, anterior posterior] pairs, with bregma = [0, 0], and coordinates to the left of, or posterior to, bregma reported as negative. Day 6, indicated with **, was the day that control regions were switched midway through the training session.

|  | [M/L, A/P]<br>(mm) | M1 | M2 | M3 | M4 | M5 | M6 | M7 |
| --- | --- | --- | --- | --- | --- | --- | --- | --- |
| Day 1 | R1 | -1.5, 1.5 | -2.5, 2 | -2, 1.5 | -2.5, 2 | -2.5, 2.5 | -2, 1.5 | -1.5, 1 |
|  | R2 | -2.5, 1.5 | -2.6, 1 | -2.2, 1 | -2.5, 0.5 | -2.5, 2 | -2.5, 2.5 | -2.5, 2 |
| Day 2 | R1 | -1.5, 1.5 | -2.5, 2 | -2, 1.5 | -2.5, 2 | -2.5, 2.5 | -2, 1.5 | -1.5, 1 |
|  | R2 | -2.5, 1.5 | -2.6, 1 | -2.2, 1 | -2.5, 0.5 | -2.5, 2 | -2.5, 2.5 | -2.5, 2 |
| Day 3 | R1 | -1.5, 1.5 | -2.5, 2 | -2, 1.5 | -2.5, 2 | -2.5, 2.5 | -2, 1.5 | -1.5, 1 |
|  | R2 | -2.5, 1.5 | -2.6, 1 | -2.2, 1 | -2.5, 0.5 | -2.5, 2 | -2.5, 2.5 | -2.5, 2 |
| Day 4 | R1 | -1.5, 1.5 | -2.5, 2 | -2, 1.5 | -1, 3 | -2.5, 2.5 | -2, 1.5 | -1.5, 1 |
|  | R2 | -2.5, 1.5 | -2.6, 1 | -2.2, 1 | -2, 1.5 | -2.5, 2 | -2.5, 2.5 | -2.5, 2 |
| Day 5 | R1 | -1.5, 1.5 | -2.5, 2.3 | -2, 1.5 | -2.5, 2 | -2.5, 2.5 | -1, 1 | -2, 1.5 |
|  | R2 | -2, 1.5 | -2.5, 1.5 | -2.2, 1 | -2, 1.5 | -2.5, 2 | -2, 1.3 | -1.5, 1.5 |
| Day 6** | R1 | -1.5, 1.5 | -2.5, 2.3 | -2.5, 2.5 | -2.5, 2 | -2.5, 2.5 | -1.5, 1 | -2, 1.5 |
|  | R2 | -2, 1.5 | -2.5, 1.5 | -2, 2 | -2, 1.5 | -2.5, 2 | -2, 1.5 | -1.5, 1.5 |
| Day 7 | R1 | -1, 1.5 | -2.5, 2.5 | -2.5, 2.5 | -2.5, 2 | -2, 2.5 | -2.5, 1.5 | -2, 1.5 |
|  | R2 | -1.2, 1.8 | -0.5, 3 | -2, 2 | -1.5, 2.5 | -1, 0.5 | -2.5, 2 | -1.5, 1 |
| Day 8 | R1 | -0.8, 2.2 | -1.7, 1.5 | -1, 1.5 | -2.5, 2 | -2, 2.5 | -2.5, 1.5 | -2, 2 |
|  | R2 | -1, 2 | -1, 2 | -1.5, 2 | -1.5, 1.5 | -1, 0.5 | -2.5, 2 | -2.5, 2 |
| Day 9 | R1 | -1.2, 0.5 | -2, 1.5 | -1.5, 3 | -2.5, 2 | -2, 2.5 | -2.5, 1.5 | -1.5, 1.5 |
|  | R2 | -2.2, 1 | -1.5, 1 | -1.5, 1.5 | -1.5, 0.5 | -1, 0.5 | -2.5, 2 | -2.5, 2 |
| Day 10 | R1 | -1.5, 1 | -2, 1.5 | -2.5, 2.2 | -2.5, 2 | -2, 1.5 | -2.5, 1.5 | -1.5, 1.5 |
|  | R2 | -1.2, 2 | -1.5, 1 | -1.5, 1.5 | -1.5, 0.5 | -1.5, 1.5 | -2.5, 2 | -1.7, 1.7 |
| Day 11 | R1 | -1.5, 1 | -2, 1.5 | -2.5, 2 | -2.5, 2 | -2, 1 | -2.5, 1.5 | -1.5, 1.5 |
|  | R2 | -1, +2 | -1.5, 1 | -1.5, 2 | -1.5, 0.5 | 2, 1 | -2.5, 2 | -1.7, 1.7 |
| Day 12 | R1 | -1.5, 1 | -2.5, 2 | -0.7, 0.5 | -2.5, 2 | -2, 1 | -2, 1.5 | -0.5, 0.5 |
|  | R2 | -1, 1.5 | -1.5, 1 | -2, 2 | -1.5, 0.5 | 2, 1 | -2, 2.5 | -1.5, 1.5 |
| Day 13 | R1 | -1.5, 1 | -1.2, 0.5 | -1, 0.5 | -1.5, 0.5 | -2, 2 | -1, 0.5 | -0.5, 0.5 |
|  | R2 | -1, 1.5 | -1.5, 2 | -2, 2 | -2.5, 2 | -1, 0.5 | -2, 1 | -1.5, 1.5 |
| Day 14 | R1 | -1.5, 0.2 | -1.2, 0.5 | -1, 0.5 | -1.5, 0.5 | -1.5, 1.5 |  |  |
|  | R2 | -2, 2 | -1.5, 2 | -2, 2 | -2.5, 2 | -2, 2 |  |  |
| Day 15 | R1 | -1.5, 0.2 | -1, 0.2 | -1, 0.5 | -1.5, 0.5 | -1.5, 1.5 |  |  |
|  | R2 | -2, 2 | -1.5, 2 | -2, 2 | -2.5, 2 | -2, 2 |  |  |
| Day 16 | R1 |  | -1, 0.2 | -1, 0.5 | -1.5, 0.5 | -1.5, 1.5 |  |  |
|  | R2 |  | -1.5, 2 | -2, 2 | -2.5, 2 | -2, 2 |  |  |
| Day 17 | R1 |  |  |  |  | -1.5, 1.5 |  |  |
|  | R2 |  |  |  |  | -2, 2 |  |  |

---

**Supplemental Video 1. Activity averaged around hits (related to Figure 1)**

Video of cortical activity averaged around hits. Time from hit (in seconds) indicated at the top left, and control regions are indicated (region 1 with a blue box, and region 2 with a green box). Animal performs task by sweeping activity laterally from region 1 towards region 2 at the time of the hit. Lick and forepaw activity are then evident as animal consumes reward.

**Supplemental Video 2. Activity averaged around hits (related to Figure 1)**

Video of cortical activity averaged around hits. Time from hit (in seconds) indicated at the top, and control regions are indicated (region 1 with a blue box, and region 2 with a green box). Activity in region 1 increases compared to region 2 at the time of the hit. Lick and forepaw activity are then evident as animal consumes reward.

**Supplemental Video 3. Discovery of successful activity patterns on a day of training where new control regions were introduced (related to Figures 1 & 2)**

(Top) Animals had to sufficiently increase activity in control region 1 (R1) relative to region 2 (R2) to bring the visual cursor to position 8 for 300 ms to receive reward. Video of continuous recordings from R1 (blue trace) and R2 (green trace) on a day of training when new control regions were introduced. If the animal failed to do this within 30 seconds, it was considered a failure trial (only one of every 3 frames is shown). Cyan vertical bars indicate trial starts; magenta vertical bars indicate a successful target hit; black vertical bars indicate the end of a failure trial. Top traces in black indicate running velocity (top trace) and lick bouts (second trace). The R1-R2 trace is indicated in the bottommost black trace.

(Middle) Reconstruction of the visual feedback cursor position at the time in the recording indicated by the vertical red bar in the top panel. The visual cursor disappeared from the screens in between trials.

(Bottom) Activity in dorsal cortex as animal performs the task. The location of control region 1 is denoted with a blue box, control region 2 with a green box.

